# Translesion DNA synthesis on pyrimidine dimers by Plant organellar DNA polymerases is metal-dependent

**DOI:** 10.64898/2026.01.26.701875

**Authors:** Noe Baruch-Torres, Joon Park, Eduardo Castro-Torres, Shigenori Iwai, Y. Whitney Yin, Luis G. Brieba

## Abstract

Ultraviolet (UV) radiation generates crosslinked DNA lesions—primarily cyclobutane pyrimidine dimers (CPDs) and [6–4] photoproducts ([6–4] PPs)—that block the progression of replicative DNA polymerases. In plants, these lesions are efficiently removed from nuclear DNA by dedicated repair pathways; however, comparable repair mechanisms are absent in plastids and mitochondria. Consequently, how plant organellar DNA polymerases (POPs) tolerate or bypass UV-induced damage has remained unclear. Here, we show that the two *Arabidopsis thaliana* organellar polymerases, AtPolIs, possess robust translesion synthesis (TLS) activity across CPDs. Although wild-type enzymes display only limited extension across [6–4] PPs, removal of their exonuclease function dramatically enhances bypass, yielding an efficiency of replication across the [6–4] PP that closely resembles that observed on an undamaged template. This establishes AtPolI as the first known replicative DNA polymerase capable of efficiently bypassing a [6–4] PP. We further demonstrate that TLS across UV photoproducts relies on three unique amino acid insertions within the AtPolI polymerase domain, as deletion of any single insertion abolishes TLS. Notably, Mn²⁺ can restore TLS activity in these variants, but only for CPD lesions. Together, these findings identify AtPolIs as the first plant organellar replicases with intrinsic [6–4] PP bypass capability and define the structural features that enable this function.

**GRAPHICAL ABSTRACT:** 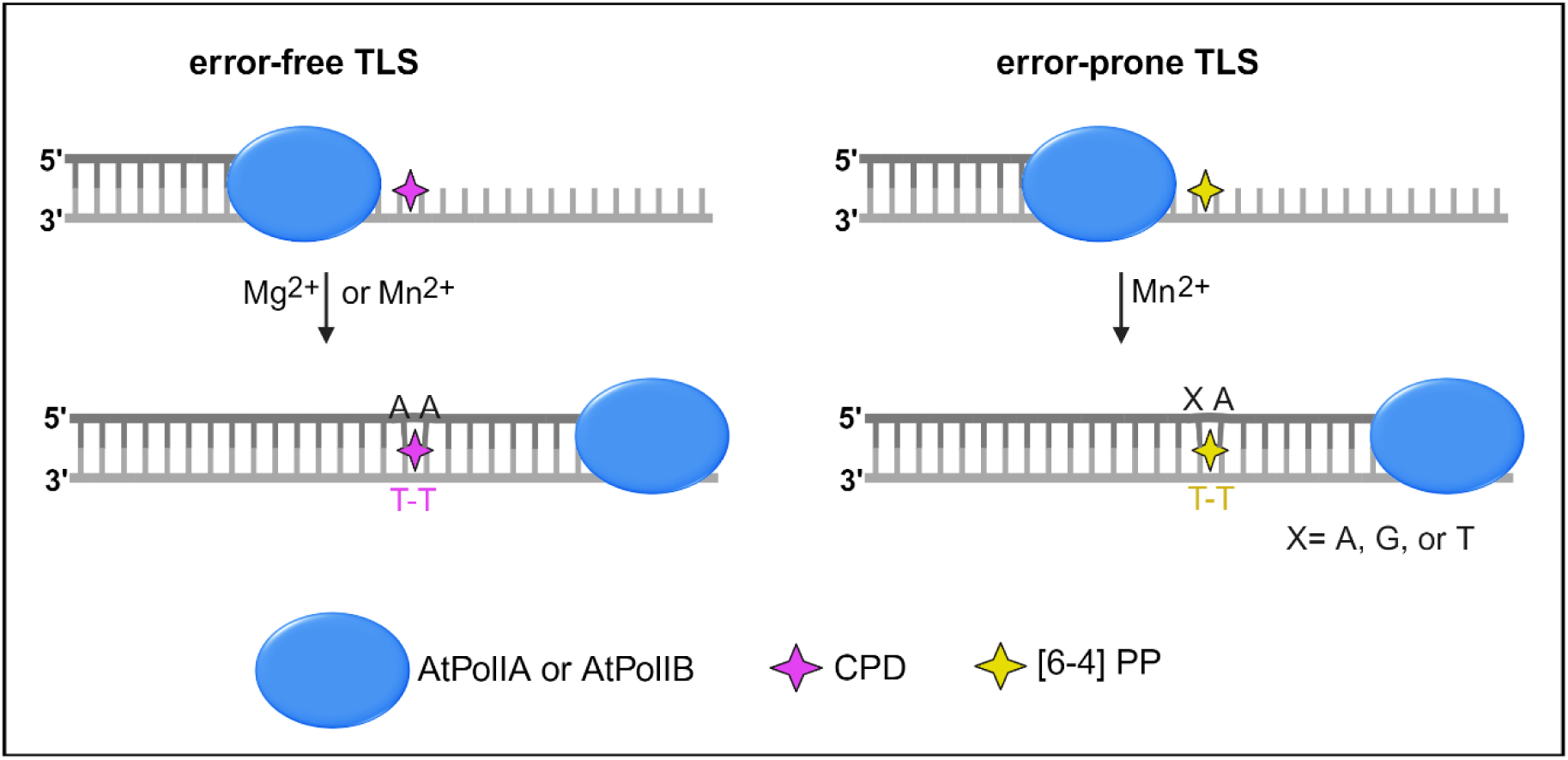

## INTRODUCTION

Ultraviolet-B (UV-B) radiation (280-315 nm), a ubiquitous DNA-damaging inducer, is especially harmful to photosynthetic organisms that are obligatory to prolonged exposure to sunlight (1, 2). UV-B promotes covalent linkages between two adjacent pyrimidines, generating cyclobutane pyrimidine dimers (CPDs) or [6-4] pyrimidine-pyrimidone photoproducts ([6-4] PPs) (3). These pyrimidine dimers can impede the progress of replicative DNA polymerases (DNAP), collapse replication fork, thus are potentially lethal to cells (4–8). Diverse routes such as fork reversal, lesion skipping, and translesion DNA synthesis (TLS) are used by the cell to cope with pyrimidine dimers during cellular replication(7, 9–11).

In contrast to the nucleus (12–15), plant organelles display an incomplete set of UV lesion repair: only photolyases but not nucleotide excision repair (NER) pathway has been detected (16–18). This suggests that DNA replication in plant organelles proceeds in the presence of CPDs and [6-4] PPs, which implies that plant organellar DNA polymerases (POPs) need to deal with UV-induced lesions during DNA replication.

*Arabidopsis thaliana* harbors two POPs (AtPolIA and AtPolIB) that share 72% of amino acid sequence identity. Although considered as organellar replicases, AtPolls also display activities of DNA repair polymerases, such as dRP lyase, microhomology-mediated end-joining (MMEJ), and translesion synthesis on abasic sites and thymine glycols (19–23). AtPolls possess a 3’-5’ exonuclease and a 5’-3’ polymerization domain. Structurally, POPs are closer related to the bacterial DNA Pol I (22, 24), Nevertheless, POPs contain three insertions in their Pol domain that resemble those in human DNA repair polymerase Pol θ (25). The POP insertions confer the ability to extend across the abasic sites and thymine glycols (20, 26).

Here we explored AtPolls’ ability to bypass CPD and [6-4] PPs lesions and evaluated the contribution of their insertion motifs to translesion synthesis activity (27, 28). We found that both AtPolIs bypass CPDs and [6-4] PPs in a metal-dependent manner. We show that AtPolIs harbor strong TLS activity on CPDs using either Mg^2+^ or Mn^2+^. While the limited [6-4] PP bypassing activity was stimulated by Mn^2+^, suppression of the exonuclease activity resulted in a drastic increase in [6-4] PP bypassing synthesis [6-4]. We also demonstrated that the POPs insertions are essential for the TLS across UV lesions.

## MATERIAL AND METHODS

### Plasmid constructs of AtPolIs

Optimized synthetic genes of *AtPolIA* (At1g50840) and *AtPolIB* (At3g20540) lacking the dual targeting sequence and the disordered region were subcloned into the *Nde* I and *BamH* I restriction sites of the modified pET19b vector containing a N-terminal 9×His tag and a PreScission protease cleavage site. AtPolIA and AtPolIB exonuclease deficient were designed as previously reported using the Q5 Site-Directed Mutagenesis protocol (New England Biolabs, Ipswich, MA, USA). Constructs for ΔIns1, ΔIns2 and ΔIns3 were designed using as background AtPolIB exo- as described in a previous report (19).

### Protein expression and purification

Plasmids containing the different AtPolIs variants were over expressed in *E. coli* Rosetta 2 (DE3) Novagen® (MilliporeSigma, Rockville, MD, USA). Bacterial cultures supplemented with 100 μgmL^-1^ of ampicillin were incubated at 37°C and 200 rpm to reach a OD_600_ of 0.5. Cultures were induced by adding 0.5 mM IPTG and incubated for 17 h at 17°C.

The different AtPolI variants were purified using the same protocol. Briefly, bacterial pellet was resuspended in a buffer containing 25 mM HEPES pH 8.0, 10% glycerol, 15 mM imidazole pH 8.0, 1 mM PMSF, 500 mM NaCl, and 0.2 mg mL^-1^ lysozyme. The resuspended solution was incubated on ice for 30 min before sonication. Lysate fraction was clarified at 30,000xg for 60 min at 4°C. The supernatant was incubated with Qiagen Ni-NTA agarose resin (Qiagen, Germantown, MD, USA) for 70 min while rocking at 4°C. The nickel resin was washed twice with lysis buffer containing 30 and 50 mM of imidazole. The protein was eluted with lysis buffer containing 500 mM imidazole and dialyzed in a D buffer (25 mM HEPES pH 8.0, 10% glycerol, 200 mM NaCl, 2 mM EDTA pH 8.0, 2 mM DTT). After dialysis, the protein fraction was concentrated and loaded to a HiTrap heparin HP column (Cytiva, Marlborough MA, USA). Bound protein was eluted using a NaCl gradient (50-1000 mM). Fractions with pure protein were pooled, concentrated and loaded into a Superdex 16/60 200 column (Cytiva, Marlborough MA, USA) and eluted using a buffer E (25 mM HEPES pH 8.0, 10% glycerol, 100 mM NaCl, 2 mM EDTA pH 8.0, 2 mM DTT). Pure fractions were pooled and concentrated in the gel filtration buffer and stored at -80°C until use.

### Oligonucleotides substrates

DNA Oligonucleotides containing a CPD (3’-TCG ATA CTG GTA CTA ATG CTT AAC GAA **T-T**A AGC ACG TCC GTA CCA TCG A-5’) and a [6-4] PP (3’- TCG ATA CTG GTA CTA ATG CTT AAC GAA **T-T**A AGC ACG TCC GTA CCA TCG A-5’) were synthetized as previously described (29, 30). An equivalent non-damaged DNA template (ND) was purchased from Integrated DNA Technologies. We purchased four FAM-labeled primers from Integrated DNA Technologies and their sequences are as follow: primer N) FAM 5’-AGC TAT GAC CAT GAT TAC GAA TTG CTT -3’, primer N+1) FAM 5’-AGC TAT GAC CAT GAT TAC GAA TTG CTT A-3’, primer N+2) FAM 5’-AGC TAT GAC CAT GAT TAC GAA TTG CTT AA-3’,primer N+3) FAM 5’-AGC TAT GAC CAT GAT TAC GAA TTG CTT AAT-3’ and a primer N+5) FAM 5’-AGC TAT GAC CAT GAT TAC GAA TTG CTT AAT TC-3’.

We annealed the following dsDNA substrates: CPD/N, CPD/N+1, CPD/N+2, CPD/N+3, [6-4] PP/N, [6-4] PP/N+1, [6-4] PP/N+2, [6-4] PP/N+3, [6-4] PP/N+5 ND/N, ND/N+1, ND/N+2, ND/N+3 and ND/N+5 by mixing the FAM-labeled primer with the complementary oligonucleotide, incubated at 95°C for 5 minutes followed by slow cooling to room temperature and stored at -20°C.

### Translesion DNA synthesis assay

We run translesion DNA synthesis assays to test AtPolIs exo+ and AtPolIs exo-variants. Reactions were performed in a buffer containing 10 mM Tris-HCl pH 7.7, 50 mM NaCl, 5% glycerol, 1.5 mM DTT, 0.2 mgmL^-1^ BSA with 100 nM of substrate ND/N, CPD/N or [6-4] PP/N mixed with 50 or 200 nM of AtPolIA and AtPolIB variants. After 5 minutes of pre incubation at 37°C, reactions were initiated by adding 200 μM of each dNTP and 2 mM of Mg^2+^ or 2 mM Mn^2+^. Reactions were quenched after 5 minutes by adding ninefolds excess of quenching buffer (80% formamide, 50 mM EDTA pH 8, 0.1% SDS, 5% glycerol, and 0.02% bromophenol blue). To test the TLS activity of the mutants ΔIns1, ΔIns2, and ΔIns3, we followed the same conditions reactions as described above.

In parallel, T7 DNAP exo+ was assayed where 100 nM of ND/N, CPD/N or [6-4] PP/N substrate and 50 or 200 nM of enzyme were combined in a buffer TR (20 mM Tris-HCl pH 7.5, 1 mM DTT, 0.2 mgmL^-1^ BSA). Reactions were incubated under the same conditions as for AtPolIs but the catalysis buffer contained 10 mM of Mg^2+^ or Mn^2+^. Samples were boiled at 95°C for 5 minutes and resolved on a 17% polyacrylamide gel containing 8M urea. Bands were visualized using FAM-fluorescence on a GE Typhoon FLA 9000 Gel Scanner (Cytiva, Marlborough, MA) and the products were quantified using ImageQuant TL (GE Healthcare). The percentage of full-length products was calculated using the ratio of full-length extension band intensity to the sum of the total band intensities in each lane.

### Exonuclease assays

To evaluate the exonuclease activity of the different AtPolls variants and T7 DNAP, we followed the same conditions for the translesion DNA synthesis assay as described above but the catalysis buffer did not contain dNTPs. Samples were boiled at 95°C for 5 minutes and resolved on a 17% polyacrylamide gel containing 8M urea. Bands were visualized using FAM-fluorescence on a GE Typhoon FLA 9000 Gel Scanner (Cytiva, Marlborough, MA).

### TLS activity under divalent metal ions titration

To characterize the influence of different Mg^2+^ and Mn^2+^ concentrations on the AtPolls activities, we performed DNA extension assays by combining 100 nM of DNA substrate (ND/PN, CPD/PN, [6-4] PP) with 200 nM of AtPolIA exo+ or AtPolIB exo+. The reactions started by adding 0.2 mM dNTPs and a gradient of 0-10 mM of Mg^2+^ or Mn^2+^. After 5 min of incubation at 37°C, sample were quenched by adding ninefolds excess of quenching buffer (80% formamide, 50 mM EDTA pH 8, 0.1% SDS, 5% glycerol, and 0.02% bromophenol blue). Quenched samples were boiled at 95°C for 5 min and resolved on a 17% polyacrylamide gel containing 8 M urea. Bands were visualized using FAM-fluorescence on a GE Typhoon FLA 9000 Gel Scanner (Cytiva, Marlborough, MA).

### DNA extension of UV lesion-containing substrates

To evaluate whether AtPolIs can extend intermediate product mimicking stages of CPD and [6-4] PP, we used the dsDNA substrates CPD/N, CPD/N+1, CPD/N+2, CPD/N+3, [6-4] PP/N, [6-4] PP/N+1, [6-4] PP/N+2 and [6-4] and PP/N+3. The 3’ OH end of the N primer is right before the lesion, N+1 covers half lesion, N+2 covers the whole lesion and N+3 pairs one base after the lesion. 100 nM of DNA and 200 nM of DNA polymerase (AtPolIA exo+, AtPolIA exo-, AtPolIB exo+, AtPolIB exo-) were mixed in 10 mM Tris-HCl pH 7.7, 50 mM NaCl, 5% glycerol, 1.5 mM DTT, 0.2 mgmL^-1^ BSA. Reactions were preincubated at 37°C for 5 minutes and started by addition of 0.2 mM of each dNTP and 2 mM of Mg^2+^ or Mn^2+^. As a control, we run reactions by combining 100 nM of the equivalent non-damaged DNA (ND) with 200 nM of DNA polymerase. Reactions were stopped at 5 min by the addition of ninefolds excess of quenching buffer (80% formamide, 50 mM EDTA pH 8, 0.1% SDS, 5% glycerol, and 0.02% bromophenol blue). Quenched samples were boiled at 95°C for 5 min and resolved on a 17% polyacrylamide gel containing 8 M urea. Bands were visualized using FAM-fluorescence on a GE Typhoon FLA 9000 Gel Scanner (Cytiva, Marlborough, MA). Bands were quantified using ImageQuant TL (GE Healthcare). The percentage of full-length product was calculated using the ratio of full-length extension band intensity to the sum of the total band intensities in each lane.

### Single nucleotide incorporation

To determine the incorporation fidelity opposite CPD and [6-4] PP, we used the dsDNA substrates CPD/N, CPD/N+1, CPD/N+2, CPD/N+3, [6-4] PP/N, [6-4] PP/N+1, [6-4] PP/N+2 and [6-4] PP/N+3, and their equivalent ND substrates. 100 nM DNA substrate and 200 nM of AtPolIA exo- or AtPolIB exo- were pre incubated at 37°C in a buffer containing 10 mM Tris-HCl pH 7.7, 50 mM NaCl, 5% glycerol, 1.5 mM DTT, 0.2 mgmL^-1^ BSA. Reactions started by the addition of 2 mM Mg^2+^ and 2.5 μM of dATP, dTTP, dGTP or dCTP (for ND), 100 μM for CPD, and 400 μM for [6-4] PP. All samples were stopped by the addition of ninefolds excess quenching buffer (80% formamide, 50 mM EDTA pH 8, 0.1% SDS, 5% glycerol, and 0.02% bromophenol blue) after 1 min at 37°C by except reactions containing [6-4] PP/PN+1 which were stopped after 5 min to ensure incorporation. Quenched samples were boiled at 95°C for 5 min and resolved on a 20% polyacrylamide gel containing 8 M urea. Bands were visualized using FAM-fluorescence on a GE Typhoon FLA 9000 Gel Scanner (Cytiva, Marlborough, MA).

## RESULTS

### AtPolIs display TLS activity across CPD and [6-4] PP

In previous works we showed that AtPolIs from *Arabidopsis thaliana* bypass DNA lesions such thymine glycols and abasic sites (19, 20). To characterize the ability of these plant organellar DNAPs to bypass more detrimental and voluminous DNA damages, we examined their TLS activity on two UV lesions: CPD and [6-4] PP in the presence of Mg^2+^ or Mn^2+^ on a substrate formed by a 49-nt DNA template with a CPD or [6-4] PP and a 27-nt primer with the 3ꞌ-OH end right before the lesion (Fig. 1A). A non-damaged DNA substrate (ND) was made with identical sequence except replacing the lesion with two thymines (Fig. 1A *bottom*). We performed the assays using sub-stoichiometric and 2-folds excess of enzyme to determine the optimal condition where TLS appears. In the presence of 2 mM Mg²⁺, AtPolIA and AtPolIB, extended 75% and 62% of the primer, respectively, across CPD on the template (Fig. 1B, E), demonstrating substantial capacity for CPD lesion bypass. In contrast, AtPolls incorporated only a single nucleotide without further extension [6-4] PP (Fig. 1F *left* and Fig. 1B *right*). Interestingly, when the assays were repeated using 2 mM Mn²⁺, the overall TLS activities were markedly enhanced. We observed the largest bypassing increment at 50 nM enzyme concentration where CPD bypassing products increased 7-folds for AtPollA and 6-folds for AtPolIB in comparison to the same conducted at 2 mM Mg^2+^ (Fig. 1C *left* and Fig. 1E *left*). On the [6-4] PP template, AtPolIA extended ⁓8% and AtPolIB ⁓11% of primers (Fig. 1C and 1F). The AtPolls extended nearly 100% of the primer to full-length on non-damaged (ND) template in the presence either Mg^2+^ or Mn^2+^ (Fig. 1D). Although the bypass efficiency was modest, these results suggest that AtPolls would be the only replicative DNA polymerase reported to perform [6-4] PP bypass and extension.

**Figure 1.**
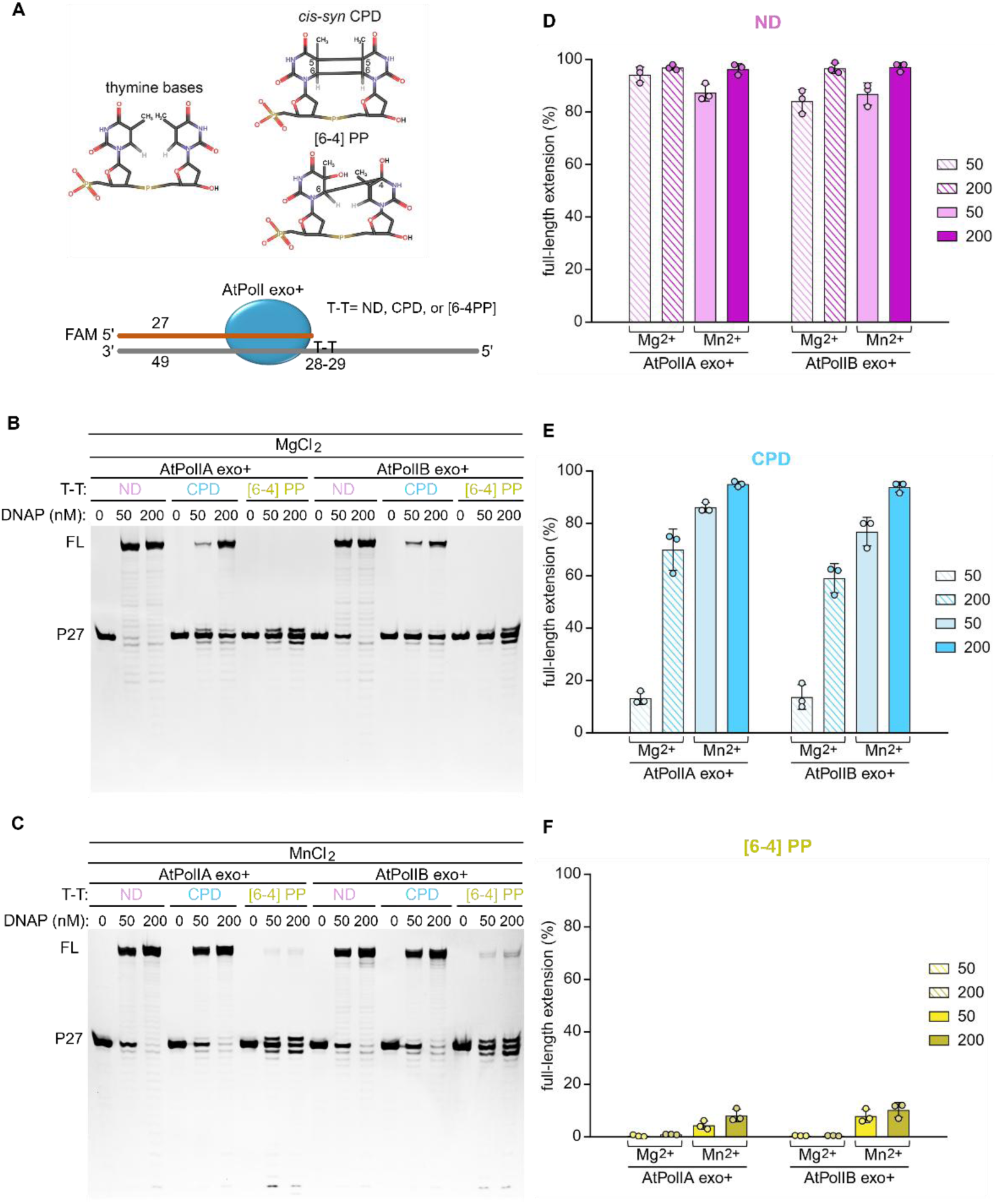
Translesion DNA synthesis assays on CPD and [6-4] PP by AtPolls exo+. (**A**) Schematic representation of a cyclobutane pyrimidine dimer (CPD) and a [6-4] PP (*top*). Cartoon depicting the DNA substrate used for lesion bypassing assays (*bottom*). (**B**) 17% denaturing gel displaying lesion bypassing assays where 50 or 200 nM of AtPolIA exo+ or AtPolIB exo+ were used and combined to 100 nM of DNA substrate (CPD, [6-4] PP, or ND) in the presence of 2.0 mM Mg^2+^. (**C**) primer extension as in panel B but using 2.0 mM Mn^2+^. (**D-F**) Plots represent the percentage of extended primer for Non damaged (magenta), CPD (cyan) and [6-4] PP (yellow). Error bars represent mean and standard deviation from at least two repeats.

We next assayed T7 DNA polymerase (T7 DNAP) TLS in the presence of Mn²⁺ to evaluate whether Mn²⁺ can stimulate lesion bypass to other DNA polymerases. No bypass products were detected on either the CPD or [6-4] PP templates (Fig. S2B). Instead, Mn²⁺ drastically reduced T7 DNAP activity on the ND template, consistent with previous reports (31) (Fig. S2C).

To evaluate whether the exonuclease activity changes across normal or damaged DNA, we examined the exonuclease activity on UV-lesion containing substrates TLS (Fig. S3A). In reactions containing the ND, CPD, and [6-4] PP with Mg^2+^, the exonucleolytic product formation ranged from 27 to 14 nts, with slightly higher activity for AtPolIA exo+ than AtPolIB exo+ (Fig. S3B). Mn^2+^ enhanced the exonuclease activity on the three substrates (ND, CPD, and [6-4] PP) following a similar trend where AtPolIB exo+ showed lower activity than AtPolIA exo+ (Fig. S3C). In contrast with AtPolIs, T7 DNAP exo+ exhibited a robust exonuclease activity in all DNA substrates with products migrating along with the running buffer in the presence of Mg^2+^ while in the presence of Mn^2+^ the activity was reduced (Fig. S3D).

These results suggest that the intrinsic TLS activity of AtPolIs to bypass UV lesion relies on its modest exonuclease activity compared with T7 DNA polymerase, whose strong exonuclease/editing function markedly reduces its bypass capacity.

### Optimal metal ion concentrations for TLS activity of AtPolIs

As AtPolIs displayed metal-dependent TLS, we investigated the optimal metal ion concentrations that support the AtPolIs TLS activity. Assays were conducted at 0.05-10.0 mM Mg^2+^ or Mn^2+^. On the ND template, the primer extension to full-length (FL) product occurred at 0.2-10 mM Mg^2+^ uniformly (Fig. 2A *left*); whereas on the CPD-containing template, the FL product appeared at 1.0 mM Mg^2+^ and reached the maximum at 2.5-5 mM Mg^2+^, where 63% of FL product was formed (Fig. 2A *middle*). As Mg^2+^ concentration increased, AtPolIA exo+ did not increase FL product synthesis but generated more exonucleolytic products, higher than those on the ND template (Fig. 2A *middle* compared to 2A *left*). On the [6-4] PP reactions, AtPollA exo+ showed no metal concentration dependency (Fig. 2A *right*).

**Figure 2.**
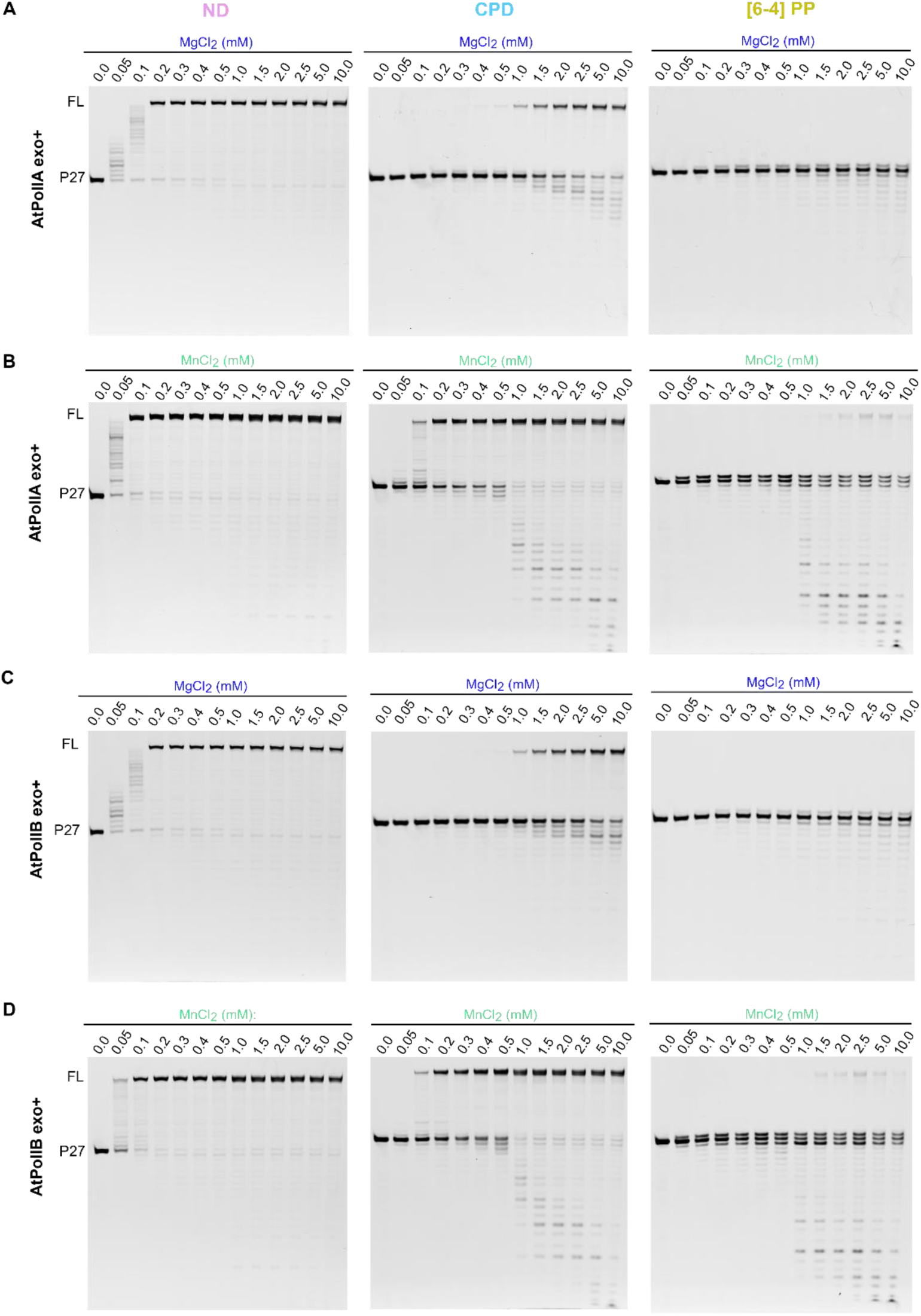
Metal-dependent translesion DNAs synthesis assays of AtPolIs exo+. (**A**) Primer extension reactions using 100 nM of DNA substrate and 200 nM of AtPolIA exo+. Primer extension of a non-damaged (*left*), CPD (*middle*) or [6-4] PP (*right*) DNA template under a 0-10 mM Mg^2+^ gradient. (**B**) as in panel A but using a gradient of Mn^2+^ from 0 to 10 mM. (**C**) As in panel A but reactions were incubated in the presence of AtPolIB exo+. (**D**) As in panel C but using a 0 to 10 mM Mn^2+^ gradient. FL, represents the extension product while the 27 band corresponds to the primer.

In the presence of Mn^2+^, AtPolIA exo+ synthesized FL product at 0.05 mM on the ND template, indicating that Mn^2+^ binds to the Pol site with higher affinity than Mg^2+^ (Fig. 2B *left* compared to 2A *left*). On the CPD template, the FL product began at 0.1 mM and reached the maximum at 0.4-0.5 mM Mn^2+^ with ⁓80% FL product, and simultaneously, an elevated exonuclease activity was observed at 1-10 mM Mn^2+^ (Fig. 2B *middle*). On the [6-4] PP template, AtPolIA exo+ extended about 11% at 2.0-5.0 mM Mn^2+^ (Fig. 2B *right*). Although the exonucleolytic pattern is similar to on the CPD template, the unextended primer in [6-4] PP reactions are not further digested (Fig. 2B *middle* compared to 2B *right*).

We repeated the same experiments using AtPolIB exo+ to compare the two homologs TLS activity on UV lesions. AtPolIB exo+ exhibited similar metal dependent activities behavior on ND, CPD and [6-4] PP as AtPolIA exo+, full primer extension at 0.2 to 10 mM Mg^2+^ on ND-template started from; 1 to 10 mM Mg^2+^ on CPD-template, and no bypassing activity on the [6-4] PP template (Fig. 2C). However, we noticed that AtPolIB exo+ produced shorter exonucleolytic products than AtPolIA exo+ on the CPD reactions (Fig. 2C *middle* compared to 2A *middle*). showing weaker exonuclease activity.

In the presence of Mn^2+^, AtPolIB exo+ showed a wider functional range of Mn^2+^ on the CPD template and generated greater than 90% FL products from 1-10 mM Mn^2+^ (Fig. 2D *middle*). On [6-4] PP reactions, Mn^2+^ triggered a limited extension of about 10% (1.5-5 mM) (Fig. 2D *right*). Overall, these results suggest a correlation between the metal ions concentration, the exonuclease, and the TLS activity.

### Extension of intermediates CPD and [6-4] PP by wild-type AtPolIs

To investigate whether exo-proficient AtPolIs can extend stalled products during CPD and [6-4] PP replication, we designed primers that mimic sequential bypassing of a UV lesion at 1-nt increment, termed N, N+1, N+2 and N+3 (Fig. 3A). As expected, both AtPolIs extended all the four ND templates with the same efficiency (Fig. 3B-C and Fig. S4B *left* and S4D *left*).

**Figure 3.**
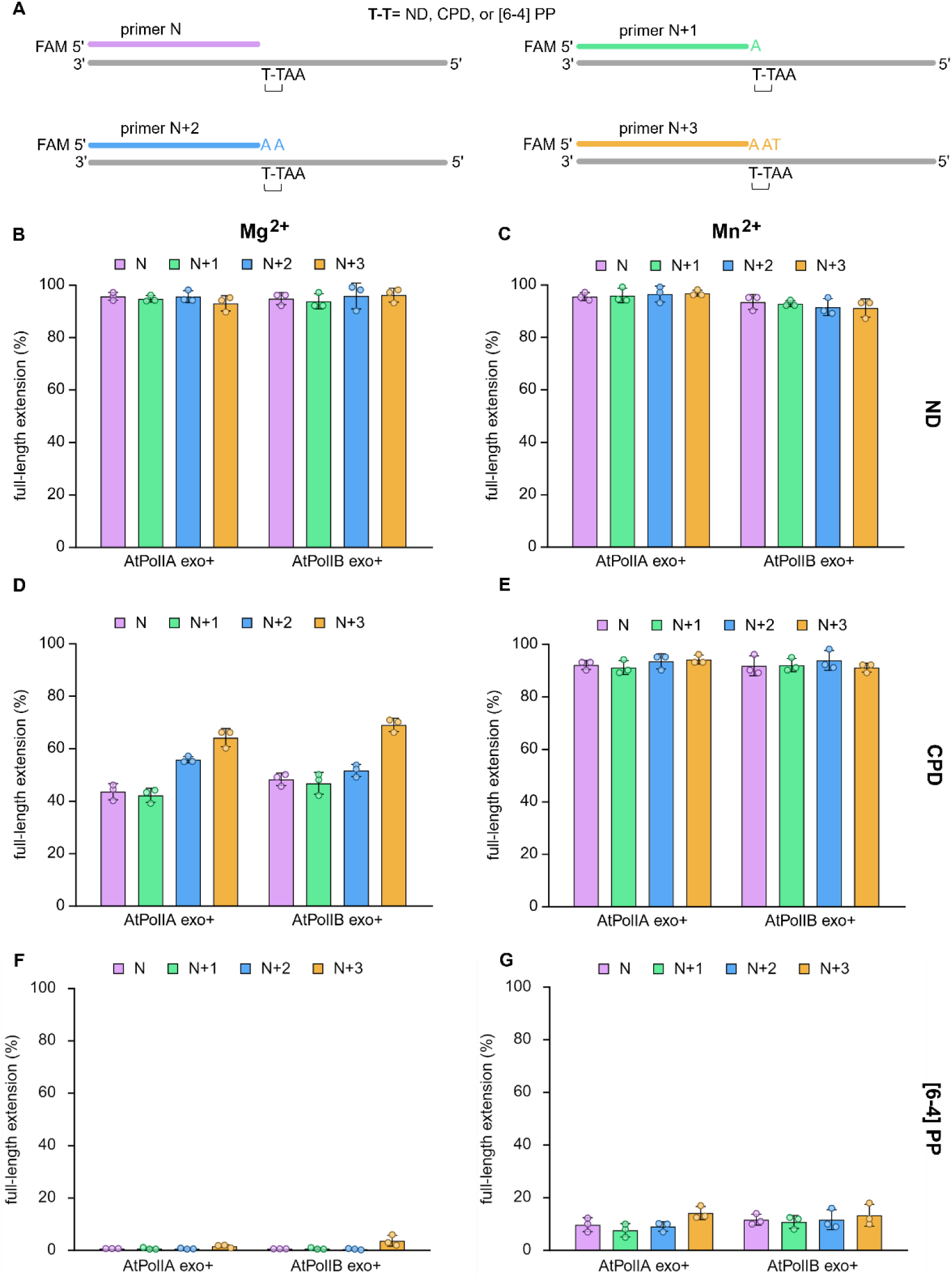
Primer extension of UV lesion-containing DNA in the presence of Mg^2+^ or Mn^2+^ by exo-proficient AtPolIs. (**A**) Scheme of the four DNA substrates used for each template (ND, CPD or [6-4] PP) where the FAM labeled primer increases by 1 nt. (**B**) Plot indicating the full extension by AtPolIA exo+ or AtPolIB exo+ incubated with four different non-damages substrates (n, N1, N2, N3, N4) in the presence of 2 mM Mg^2+^ (**B**) or 2mM Mn^2+^ (**C**). Extension of four CPD substrates by AtPolIA exo or AtPolIB exo+ using 2 mM Mg^2+^ (**D**) or 2 Mm Mn^2+^ (**E**). As in D-E but using four substrates containing a [6-4] PP lesion in the presence of 2 mM Mg^2+^ (**F**) or 2 mM Mn^2+^ (**G**).

On the CPD template and with 2 mM Mg^2+^, AtPolIA exo+ extended ⁓45% of the N and N+1 primer, 57% of N+2, and 65% of N+3 primer to FL product (Fig. 3D and S4B *middle*). Nevertheless, in the presence of 2 mM Mn^2+^, 90% of all four primers were extended to FL products (Fig. 3E and Fig. S4D *middle*).

AtPoIIB exo+ extended comparable quantity of primer across the CPD as AtPolIA exo+, displaying the highest FL product with primer N+3 whose 3ꞌ-terminus located one base after the lesion (Fig. 3D and Fig. S4C *middle*). As observed for AtPolIA exo+, AtPolIB exo+ with 2 mM Mn^2+^ extended all four primers by about 90% across CPD (Fig. 3E, and Fig. S4D *middle*). Overall, the results suggest that the primer length is positively correlated with the ability of AtPolIs exo+ trans-CPD lesion extension in the presence of Mg^2+^, whereas Mn^2+^ stimulates all primer trans-CPD extension uniformly.

On the [6-4] PP template, neither AtPolIs were able to extend any primer across the lesion with 2 mM Mg^2+^ (Fig. 3F and Fig. S4C *right*). In the presence of 2 mM Mn^2+^, AtPolIA exo+ extended 8-14% primers N, N+1, N+2, N+3, and AtPolIB exo+ 10-13% to FL product (Fig. 3G), which is modestly increased in comparison to reactions with Mg^2+^ (Fig. 3F compared to Fig. 3G).

One interesting observation is that in reaction containing Mg^2+^, the primer leftover for all CPD and [6-4] PP substrates (N, N+1, N+2, N+3) resemble the N primer length (27nt) instead of 28, 29, 30 nt for N+1, N+2, N+3 primers, respectively (Fig. S4B and S4C). These suggest that AtPolIs exo+ still detects the UV lesion even one position after the lesion (N+3), starting an excision activity that preferentially terminates when the 3’-OH primer is located right before the lesion. The results suggest that Plant Organellar Polymerases generated at least two groups of extended primers, one group treadmills at the lesion site where one nucleotide incorporation and excision occur alternatively, and another group of primer is fully extended. Interestingly, AtPolIB exo+ displays a less efficient cleaving ability on the [6-4] PP templates consistent with its lower exonuclease activity in comparison to AtPolI A exo+ (Fig. S4C *right*).

To address this lesion-detection ability, we increased the primer length to 32-nt primer (N+5) where the 3’-end primer is 3-nt post to the [6-4] PP lesion on the template (Fig. 4A). The 6-4 PP bypassing products by both AtPolIs exo+ are indeed increased to 46-50% (Fig. 4B-C). These values represent a 25-folds and 12-folds increase in [6-4] PP extension by AtPolIA exo+ and AtPolIB exo+, respectively, relative primer N+3 (Fig. 4C). The findings support the idea that the further the 3’-primer from the lesion, the lower the AtPolIs lesion detection capability.

**Figure 4.**
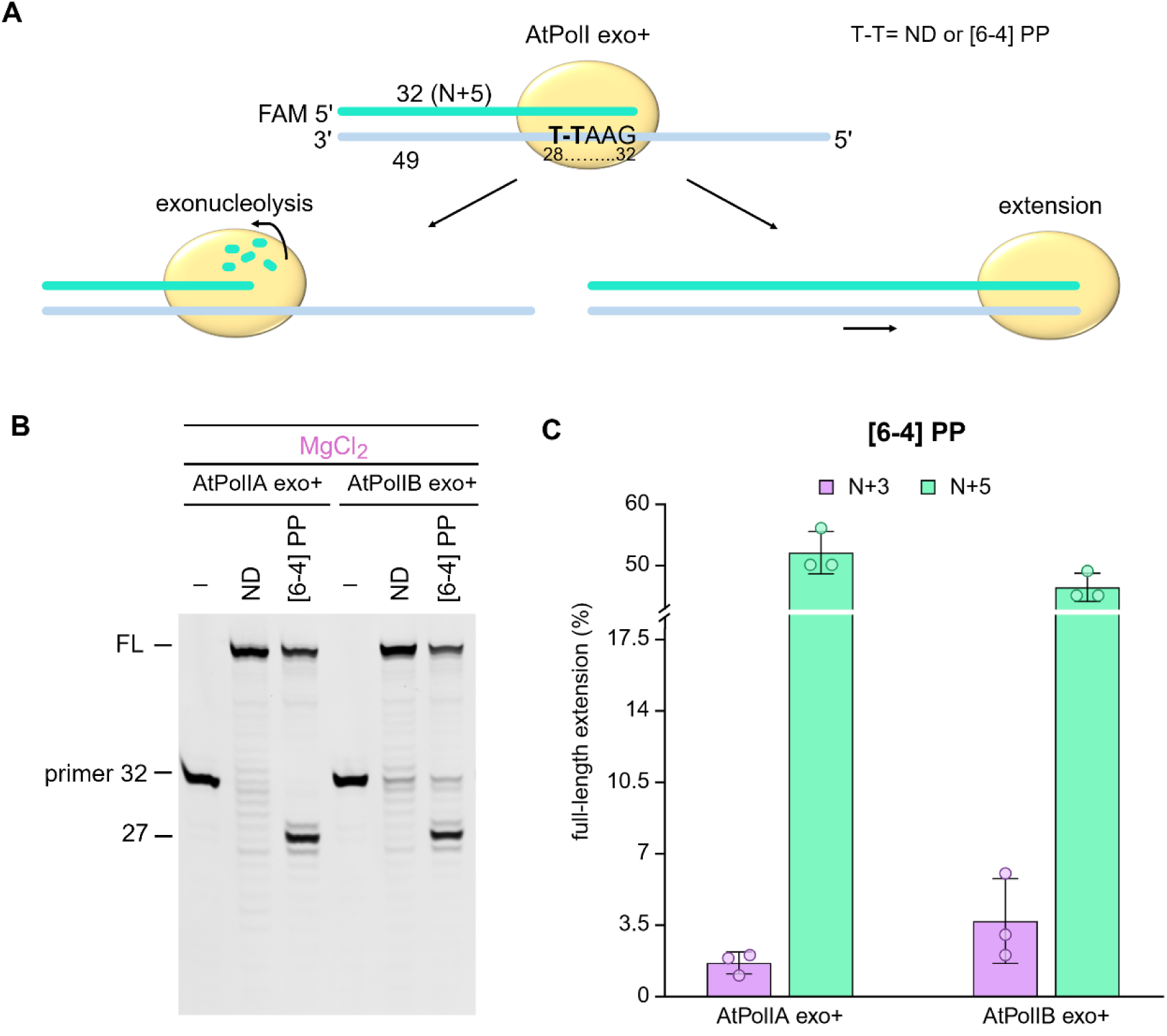
[6-4] PP extension by AtPolIs exo+ in the presence of Mg^2+^. (**A**) Cartoon of the DNA substrate where the primer is located +3 position after the lesion. The scheme shows the two possible mechanisms when AtPolIs exo+ detect a bypassed lesion. (**B**) Extension assay of a non-damaged (ND) or [6-4] PP template in the presence of 2 mM Mg^2+^. FL indicates full-length product; primer substrate is shown as a 32-nt band and the exonucleolytic product as 27-nt (right before the [6-4] PP lesion position). (**C**) Plot indicating the full-length extension by AtPolIA exo and AtPolIB exo+ on the [6-4] PP/N+3 (replotted from Fig. 3F) and [6-4] PP/N+5 (from panel B) in the presence of 2 mM Mg^2+^.

### Mn^2+^ enables the exo-deficient AtPolIs robust synthesis across [6-4] PP

We previously demonstrated that AtPolIs display moderate exonuclease activity resulting in a lower fidelity, compared to human mitochondrial replicase Pol γ and T7 DNAP, as a trade-off for their ability to bypass DNA lesions (19, 32). As the inactivation of the exonuclease domain in some DNAPs potentiate their intrinsic TLS activities (19, 33–35), we studied the TLS activity on UV lesions by AtPolIs exo-variants. We used the same CPD, [6-4] PP and a non-damaged template used for the AtPolIs exo+ (Fig. 5A). On the non-damaged template (ND), AtPolIA exo- and AtPolIB exo- completely synthesized the substrate, displaying over 95% of product in reaction containing either 2 mM of Mg^2+^ or 2 mM Mn^2+^ (Fig. 5B-D). On the CPD template, 65-82% of the primer in the presence of 2 mM Mg^2+^, and over 95% of primer in the presence of Mn^2+^ were extended by AtPolIB exo- to FL product (Fig. 5B-C and 5E). In reaction containing 2 mM Mg^2+^, AtPolIA exo- extended from 77-82% of the CPD substrate when incubated with 2 mM Mg^2+^ and over 95% in reaction using 2 mM Mn^2+^ (Fig. 5B and 5E). On the [6-4] PP substrate and with 2 mM Mg^2+^, both exo-deficient AtPolIs were unable to generate extension products (Fig. 5B and 5F). In the presence of 2 mM Mn^2+^, PolIB exo-, was able to extend 57% and 76% while AtPolIA exo- reached 62% and 80% of [6-4] PP extension (Fig. 5C and 5F). Overall, diminishing the exonuclease activity enhanced the TLS activity on AtPolIs exo-deficient variants. AtPolIB exo- increased up to 4-folds its lesion bypassing ability relative to its exo-proficient variant, while AtPolIA exo- displayed 5-folds of increment (Fig. 1E compared to Fig. 5E). In [6-4] PP reactions containing 2 mM Mn^2+^, AtPolIB exo- enhanced the extension efficiency by 7-folds while AtPolIA exo-displayed up to 10-folds product increment (Fig. 1F compared to 5F).

**Figure 5.**
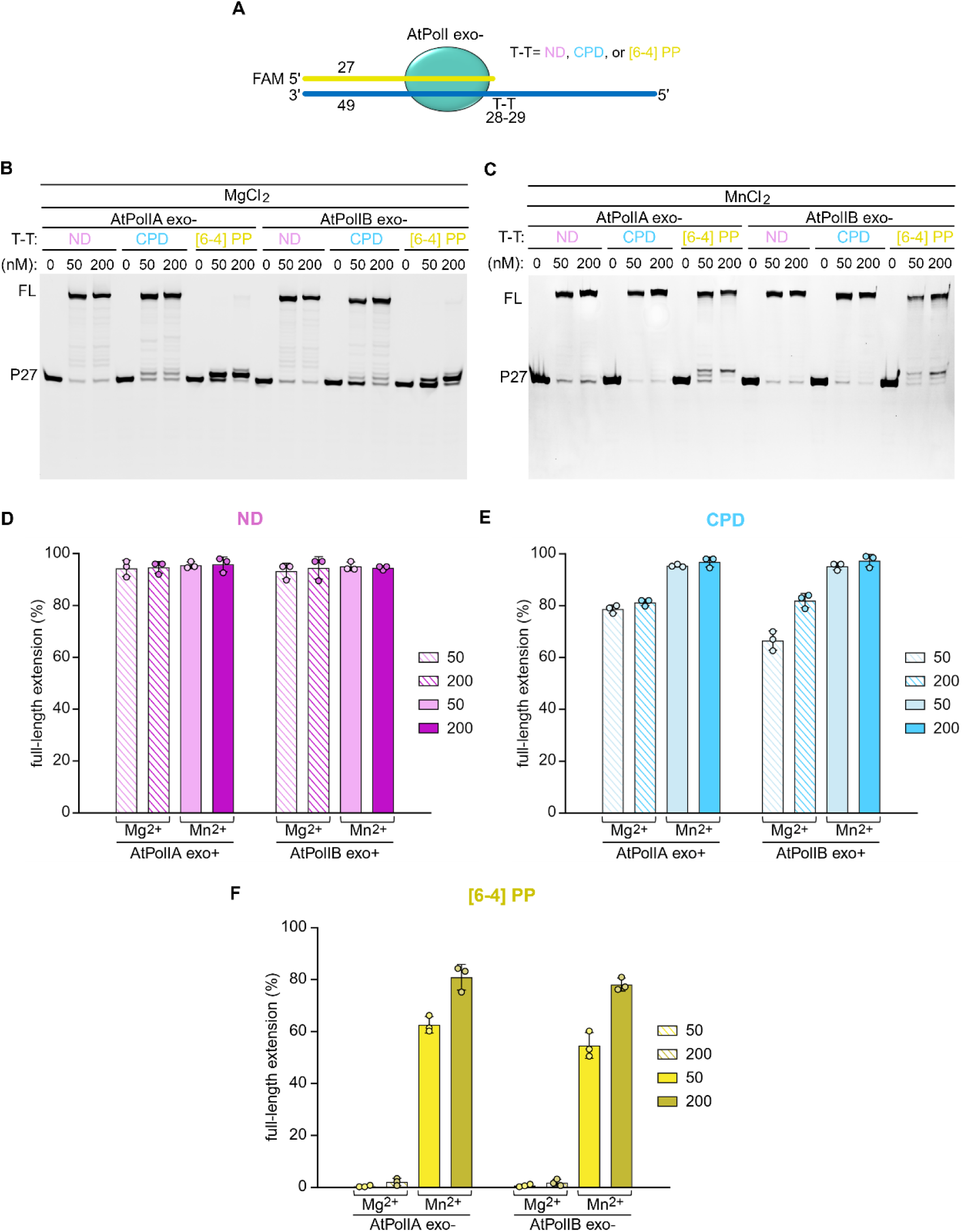
Lesion bypassing assays on CPD and [6-4] PP by exo-deficient AtPolIs. (**A**) DNA substrates used in this assay constituted by a 27-nt FAM labeled primer and a 49-nt template T-T denotes a CPD, [6-4] PP or two adjacent thymine.(**B**) Lesion bypassing assays where 50 or 200 nM of AtPolIA exo- or AtPolIB exo- were combined with 100 nM of DNA substrate in the presence of 2.0 mM Mg^2+^. (**C**) Primer extension as in panel B but using 2.0 mM Mn^2+^. Bar chart from panel B and C, indicating the percentage of extended primer for Non damaged (**D**), CPD (**E**) and [6-4] PP (**F**) substrates. Error bars represent mean and standard deviation from at least two repeats.

### Exo-deficient AtPolIs extend [6-4] PP DNA intermediates with high efficiency

We performed AtPolIs exo- primer extension assays using primers mimicking progressive bypassing UV lesion. On the CPD template, AtPolIB exo- extended 90% of primers N, N1, N+2 and N+3 to FL in the presence of Mg^2+^, and greater than 95% in the presence of 2 mM Mn^2+^ (Fig. 6D-E, S5C-E). While AtPolIB exo- primer extension on ND template with similar efficiency in the presence of either metal ions (Fig. 6A-C and S5C-E), its TLS activity has drastically increased relative to its exo+ counterpart. Similar observation was made with AtPolIA exo- (Fig. 6B-E and S5B and S5D).

**Figure 6.**
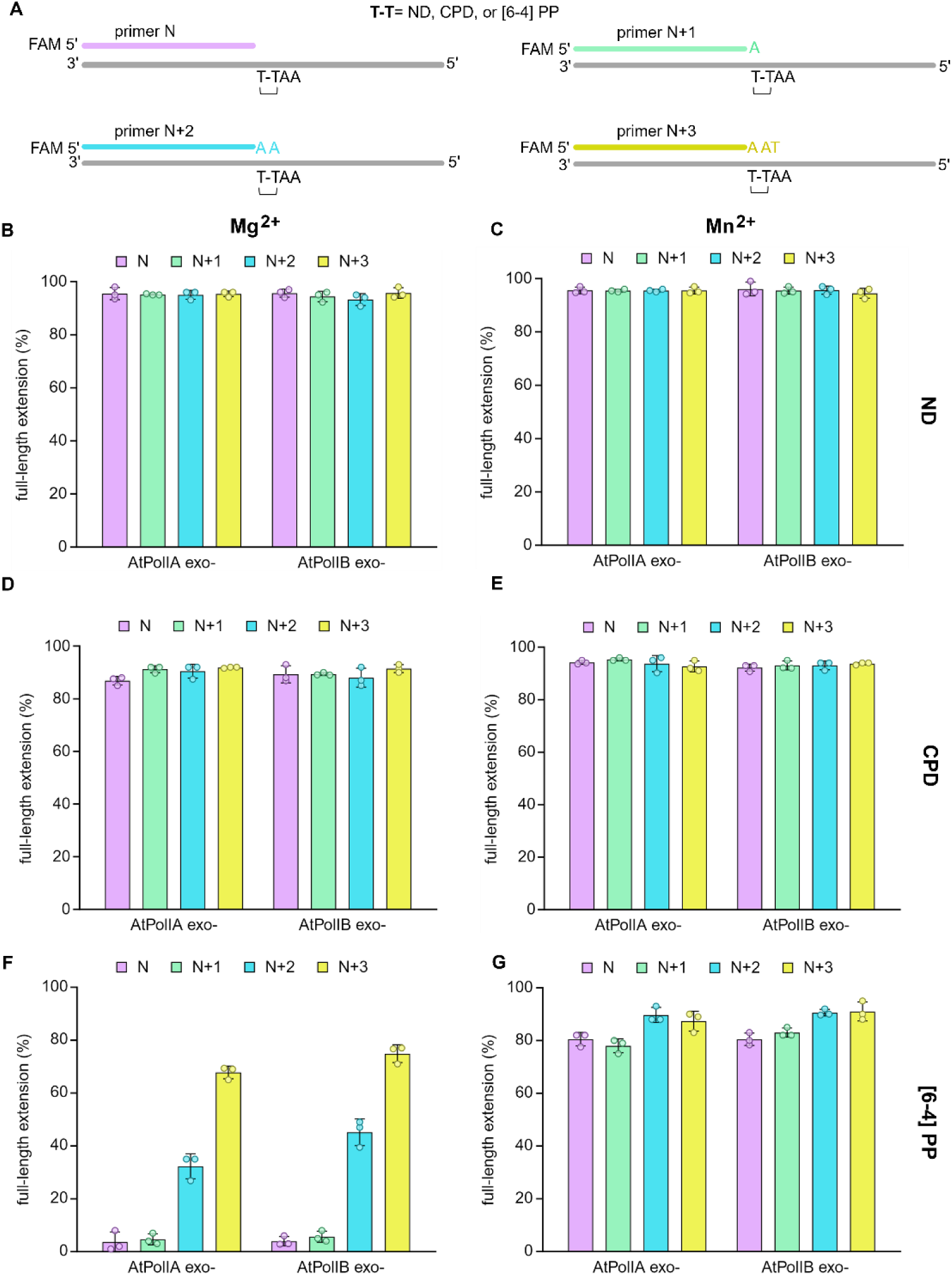
UV lesion extensions in the presence of Mg^2+^ or Mn^2+^ by exo-deficient AtPolIs. (**A**) Scheme of the twelve DNA substrates used in this assay. FAM labeled primer varies by 1nt: 27 (N), 28 (N+1), 29 (N+2) and 30 (N+3). T-T denotes two thymines, CPD or [6-4] PP. Data from primer extension assays by AtPolIs exo- variants in the presence of 2.0 mM Mg^2+^ (**B**) or 2.0 mM Mn^2+^ (**C**). Data from bypassing assays using CPD substrates incubated with 2 mM Mg^2+^ (**D**) or 2 mM Mn^2+^ (**E**). AtPolIA exo- and AtPolIB exo- display increasing product formation from N to N+3, ranging from 5 to 70% in the presence of 2 mM Mg^2+^ (**F**) while reaction incubated with 2.0 mM Mn^2+^ drastically increased the full extension products (**G**). Error bars represent mean and standard deviation from at least two repeats.

Importantly, AtPolIB exo- bypassing [6-4] PP lesion was modestly increased in the Mg^2+^ presence (Fig. 6F and S5C), but drastically increased with Mn^2+^, where AtPolIB exo- extended 80% of primer N, 83% of primer N+1, 90% of primer N+2 and 91% of primer N+3 to FL product (Fig. 6F compared to 6G and S5E). AtPolIA exo- was also able to extend the four [6-4] PP substrates in the presence of Mg^2+^, ranging from 5 to 70% (N to N3 primers) while the incubation with 2 mM Mn^2+^ improved the extension ability up to 90% with no significant improvements across the four primers (Fig. 6F and 6G and S5B and S5D). These results suggest that exonuclease activity play a key role regulating AtPolIs translesion synthesis across CPD or [6-4] PP damage.

### Nucleotide incorporation preference of AtPolIs on CPD and [6-4] PP

In the intermediates extension assay, we found that both AtPolIs exo- failed to extend across [6-4] PP in the presence of 2 mM Mg^2+^, but could insert a single nucleotide (+1) opposite the 3’-T of the [6-4] PP (Fig. S5B-C), suggesting that incorporation of the second thymine opposite the 5’-T of the [6-4] PP constitutes an energy barrier for the primer extension. To explore whether misincorporation at the +1 position caused primer extension failure at the +2 position, we performed AtPolIs exo- single nucleotide incorporations of primers N, N+1, N+2, N+3 opposite to the [6-4] PP and CPD containing templates (Fig. 7A).

**Figure 7.**
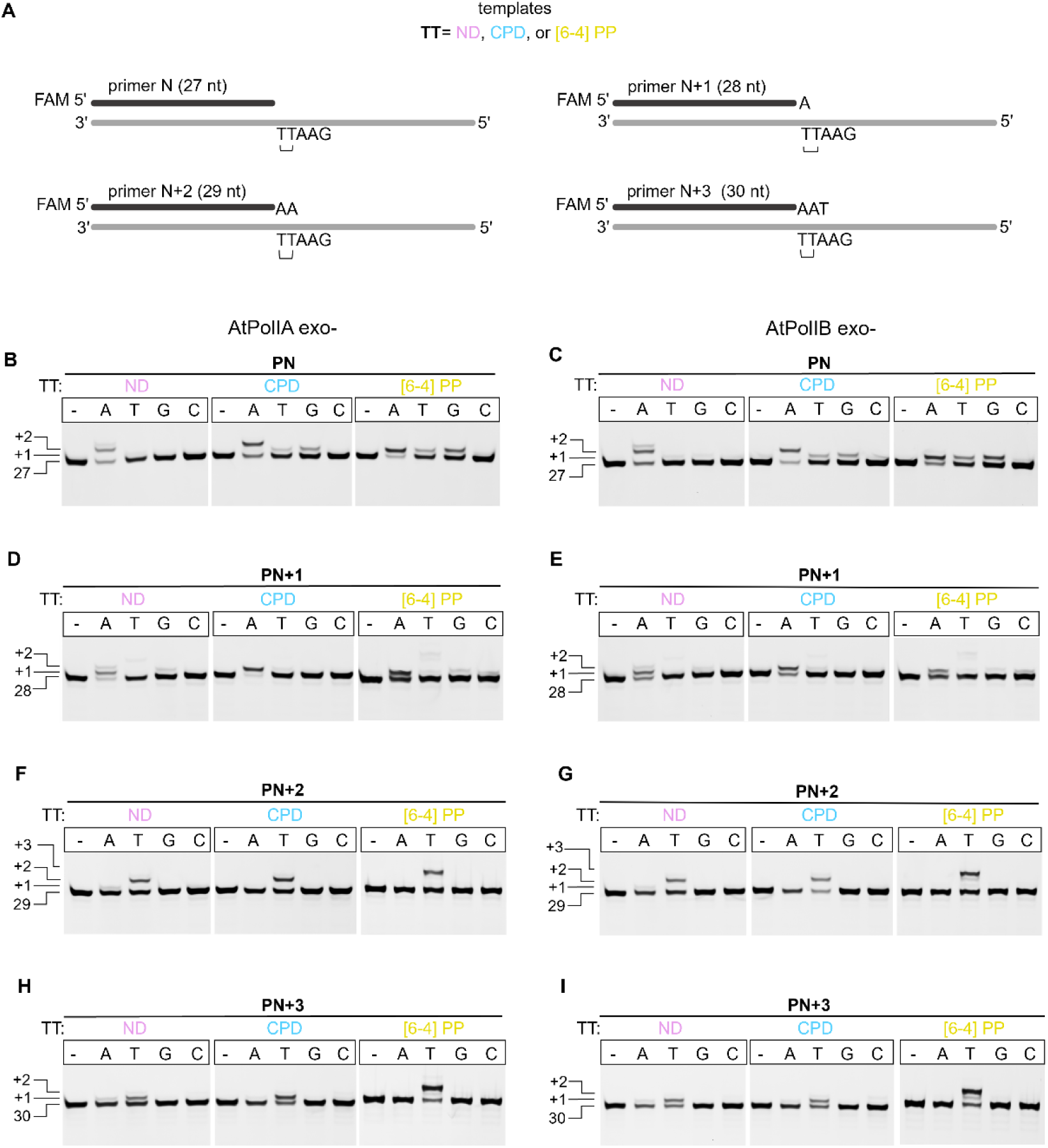
Single nucleotide incorporation catalyzed by exo-deficient AtPolIs opposite UV lesions-containing templates. (**A**) Schematic representation of the substrates used to indicate the increasing length of FAM labeled primers annealed to ND, CPD, or [6-4] PP templates. T-T denotes the location of the lesions. (**B**-**C**) Nucleotide incorporation by AtPolIA exo- and AtPolIB exo- on a ND (*left*), CPD (*middle*), and [6-4] PP (*right*) DNA template which are paired to the N primer (20nt). (**D**-**E**) individual dNTP incorporation opposite a ND, CPD, and [6-4] PP with a N+1 primer covering the first T of the lesion. (**F**-**G**) single nucleotide using a primer N+2 containing two adenines which pair to the full ring of CPD and [6-4] PP. (**H**-**I**) incorporation opposite ND, CPD, and [6-4] PP templates where the FAM labeled primer N+3 is located 1 position after the lesion. Reactions contain 2.0 mM of Mg^2+^ and 2.5 μM, 100 μM or 400 μM of each dNTP for ND, CPD and [6-4] PP, respectively. (-) denotes a reaction without dNTP. Primer and incorporation lengths are indicated on the left of the image.

Despite lack of exo activity, both AtPolIs exo- on the ND template incorporated the correct nucleotides dTMP (Fig. 7B-I *left* and Table S1-8 *left*). On the CPD substrates, AtPolIs exo- incorporated primarily correct nucleotides opposite to the +1, +2, +3 and +4 positions on the CPD template (Fig. 7B-I *middle* and Table S1-8 *middle*), with only minor misincorporation was observed at +1 position (FIG. 7B-C *middle*).

On the [6-4] PP template, AtPolIs exo- could only insert a single dAMP to the N primer opposite to the lesion and was unable to synthesize opposite to +2 position. Instead, we observed significant misincorporation at the +1 position, such as T:T and T:G mismatches (Fig. 7B-C right and Table S1-2 right)”.

We paired the first T of 6-4 PP with the N+1 primer to evaluate the incorporation at +2 position. Both AtPolIs exo- incorporated the correct an dAMP with minor dTMP, dGMP and dCMP misincorporations which generated T:T, T:G and T:C mismatches at +2 and +4 positions (Fig. 7D-E right and Table S3-4 right). With the N+2 primer, both AtPolIs exo- incorporated the two correct dTMPs at +3 and +4 positions. But an additional dTMP was mis inserted at +5 position resulting in a T:G mismatch (Fig. 7F-G right and Table S5-6 right). For the N+3 primer, AtPolIs exo- inserted the correct dTMP at +4 position but also mis incorporated a dTMP opposite the G-template (+5 position) (Fig. 7H-I right and Table S7-8 right). These results suggest that both AtPolIs execute an error-free TLS on CPD while on a [6-4] PP follows an error-prone TLS pathway.

### TLS on AtPolIs reside on unique insertions at the polymerization domain

In previous works we reported that AtPolIB amino acid insertions are involved in DNA repair and maintenance (19, 21, 23), here we aimed to evaluate the role of these insertions in AtPolIB to bypass CPD and [6-4] PP. We constructed AtPolIB exo- individual deletion mutants (ΔIns1, ΔIns2, and ΔIns3) (Fig. 8A) and tested their DNA synthesis on the ND, CPD and [6-4] PP templates (Fig. 8B). As we showed in previous assays our control AtPolIB exo- displayed full-length extension for ND, CPD but only able to insert a nucleotide opposite the [6-4] PP in reactions containing 2 mM of Mg^2+^ (Fig. 8C).

**Figure 8.**
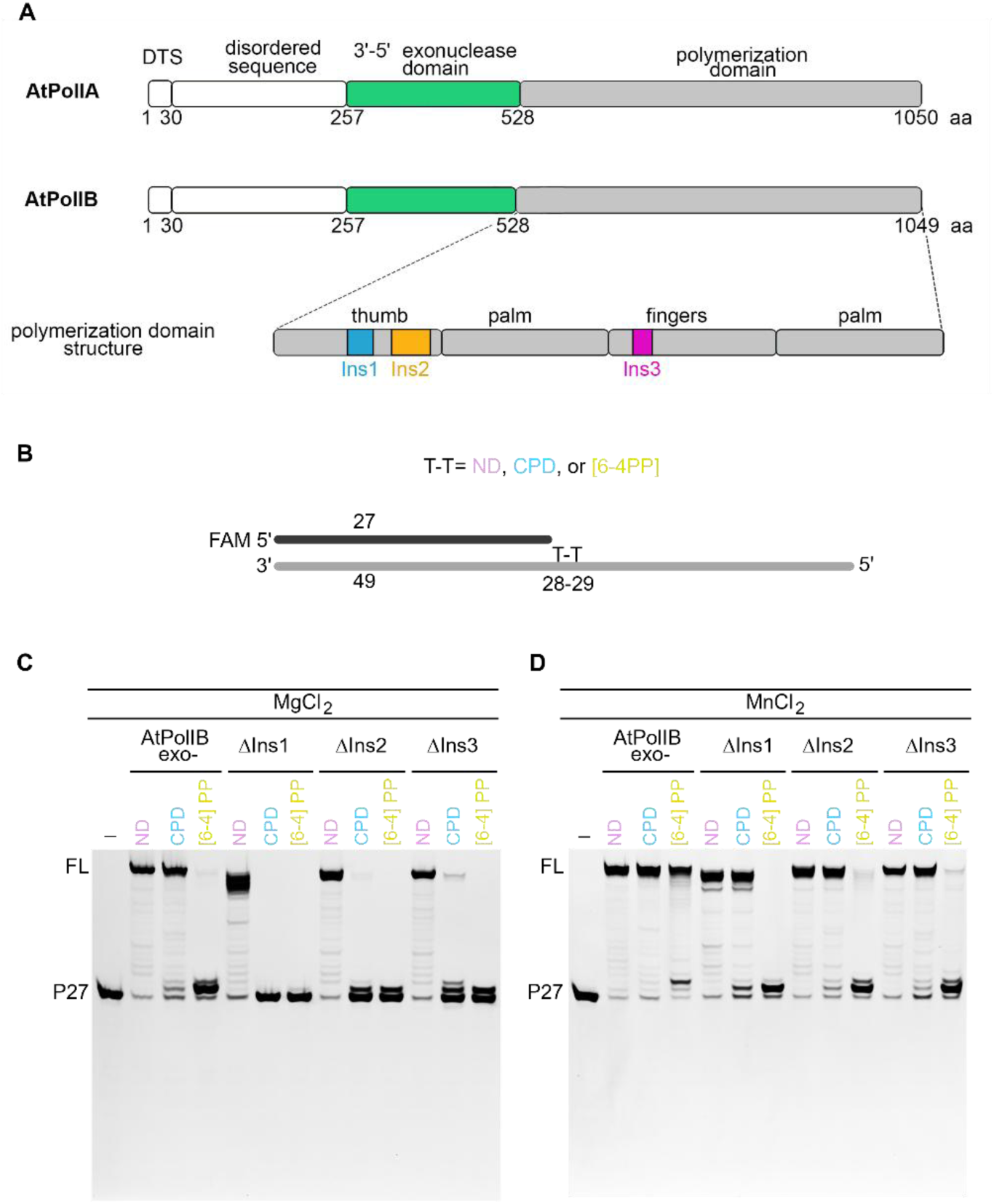
Specific aminoacid insertions in AtPolIs are involved in translesion DNA synthesis. (**A**) Schematic representation of the structural regions of AtPolIs which include a dual targeting sequence (DTS), a disordered region in the N-terminal region, a functional 3’-5’ exonuclease domain and the polymerization domain where Insertion 1, Insertion 2 and Insertion 3 are colored in navy, orange and purple, respectively. (**B**) cartoon depicting the DNA substrate where a FAM labeled 27-nt primer is annealed to a ND, CPD or [6-4] PP template. Lesion bypassing assay by AtPolIB exo-, ΔIns1, ΔIns2 and ΔIns3 in the presence of 200 μM of dNTPs and 2.0 mM of Mg^2+^ (**C**)or Mn^2+^ (**D**). In reactions incubated with Mg^2+^, all mutants and AtPolIB exo- shows full extension for ND template, AtPolIB exo- displays DNA extension on CPD while ΔIns3 shows a very reduced full extension product. In assays containing 2 mM of Mn^2+^, AtPolIB exo- generated full-length bands for ND, CPD, and [6-4] PP; ΔIns1 extended ND and CPD templates but only incorporated opposite the [6-4] PP; ΔIns2 and ΔIns3 synthesized ND, CPD and a limited extension on the [6-4] PP substrate. FL denotes full-length product.

In the presence of Mg^2+^, all deletion variants retained DNA synthesis activity on the ND template, indicating the insertions are not essential for nucleotide incorporation on nondamaged template although ΔIns1 generated shorter full-length products than in the rest of variants (Fig. 8C). Nonetheless, the deletion variants lost their translesion synthesis, the ΔIns1 and ΔIns2 variants were unable to extend across the CPD or [6-4] PP while ΔIns3 only retained a limited ability to extend CPD (Fig. 8C). Despite lacking a robust lesion extension, ΔIns2 and ΔIns3 incorporated a nucleotide opposite the first thymine on CPD or [6-4] PP while this band is absent in ΔIns1 reactions. Next, we repeated the primer extension assays in the presence of 2 mM Mn^2+^. Surprisingly, all three deletion variants regained TLS activity on the CPD template (Fig. 8D). In the presence of Mn^2+^, ΔIns1 and ΔIns3 displayed a limited TLS activity on [6-4] PP while ΔIns1 generated a single nucleotide incorporation (Fig. 8D). These results suggest that although the insertions are nonessential for synthesis on the ND template, in the presence of Mg^2+^, they are necessary for synthesis across CPD. However, in the presence of Mn^2+^, the insertions are only essential for extending [6-4] PP but not CPD.

## DISCUSSION

Plants are continuously exposed to sunlight and ultraviolet (UV) radiation that can damage DNA. To date, no replicase has been shown to possess robust lesion-bypass activity. In the nucleus, this problem is mitigated by specialized translesion DNA synthesis (TLS) polymerases and UV-damage repair pathways that bypass or remove UV lesions (44–46). However, neither TLS polymerases nor canonical repair systems are present in plant organelles, leaving organellar DNA polymerases (POPs) unavoidably confronted with UV-induced lesions during DNA replication. Nevertheless, POPs activity on UV-damaged DNA templates remains unexplored. We report that the plant *Arabidopsis thaliana* organellar DNA polymerases, AtPolIA and AtPolIB, are the first replicative DNA polymerases shown to possess intrinsic translesion synthesis (TLS) activity across UV-induced lesions. AtPolIs exhibit robust synthesis across cyclobutane pyrimidine dimers (CPDs) and limited activity across (6–4) photoproducts. Because canonical TLS polymerases lack 3’→5’ exonuclease activity, we inactivated the exonuclease function of AtPolIs and observed a drastic increase in their TLS efficiency. Together with our previous findings, these results demonstrate that AtPolIA and AtPolIB can perform translesion DNA synthesis across abasic sites, thymine glycols, and UV-induced adducts.

Family-A DNAPs exhibit different TLS abilities and efficiencies to deal with UV lesions. For instance Human Pol θ extends CPD-containing templates which would be due a lack of the 3’-5’ exonuclease activity (36); the replicative T7 DNAP is unable to replicate on CPD or [6-4] PP lesion either in the presence of Mg^2+^ or Mn^2+^ (37); *E*. *coli* DNAP I display a moderated bypassing ability on CPD but is blocked by [6-4] PP (38) while the mitochondrial replicative human Pol γ bypasses CPDs only in the presence of Mn^2+^ but it is unable to extend a [6-4] PP with either Mg^2+^ or Mn^2+^ (35).This evidence would suggest a novel mechanism that AtPolIs use when a [6-4] PP or CPD is encountered switching from a normal to a translesion DNA synthesis. In nucleus [6-4] PP bypass requires the coordinated action of two TLS DNA polymerases: the inserter and extender DNAP (39–43). Our results suggest that plant organellar DNA polymerases represent the first evidence of a replicative DNAP equipped with these abilities and extend [6-4] PP extension and modulated by Mn^2+^. These findings correlate with the fact that POPs are phylogenetically related to TLS DNAPs Pol θ and Pol ν and not with the metazoan replicative mitochondrial DNAP nor with T-odd bacteriophages (36, 44, 45) (Fig. S1). Combining our finding we propose that AtPolls, and particularly AtPolIB, have evolved four features to achieve TLS across from CPD and [6-4] PP lesions: 1) intrinsic promiscuous polymerase active site, 2) presence of low exonucleolytic or primer editing activity, 3) a strong binding affinity for the primer-template, and 4) use of Mn^2+^ that modulates the polymerase active site and key regions such as the three unique insertions found in POPs to accommodate UV-derived DNA lesion.

### 1. AtPolIs presents more promiscuous polymerase active site than conventional replicases

An intrinsic feature of TLS polymerases is their low fidelity for nucleotide incorporation. The assembly of a high-fidelity and snug active site during nucleotide incorporation in family-A DNAPs (46, 47) is challenged by *Helicobacter pylori*, *Entamoeba histolytica* DNAPI (48, 49), human Pol θ (50) and Pol ν (51) that bypass abasic sites and thymine glycol while execute nucleotide incorporation with low fidelity. AtPolIs present contrasting fidelity for nucleotide incorporation. AtPolIs exhibit near 20-folds lower fidelity for nucleotide incorporation than human counterpart, Pol γ (32). Human Pol ν exhibits low fidelity due to specific amino acids of the α-helix O, however those residues are not conserved in AtPolIs (45, 52). Human Pol ν and Pol θ harbor a specific insertion that enlarges the polymerase active site and creates a flexible active site crucial for voluminous DNA lesion accommodation in the polymerization site (52) (Fig. S1B-C). Members of the family-Y, TLS DNA polymerases share a widened active site, for instance in Pol η the CPD is perfectly accommodated in the Pol site (53). Plant organellar DNAPs seem to have evolved features found in replicative and TLS DNAPs. Although both AtPolIs share about 75% amino acid identity, AtPolIB has been associated with DNA repair and maintenance consistent with its low fidelity during nucleotide incorporation relative to AtPolIA, which is considered the replicative DNAP in plant organelles (32, 54).

The unique ability of AtPolIs to bypass UV damages would be a result of novel features to accommodate bulky lesions. AtPolIs, as other POPs, contain three unique insertions in the polymerization domain found in other DNAP such Pol θ or Pol ν but absent in bacterial DNAP I (Fig. S1). Insertion 1 is mapped from residues 576 to 621, Insertion 2 mapped from residues 648 to 712 of AtPolIB and Insertion 3 from residues 844 to 869 (Fig. S1D). When we deleted individual insertions, the TLS activity on both CPD and [6-4] PP disappeared while the presence only rescued the TLS on CPD suggesting that these insertions may influence the active site dynamics creating a more permissive active site or generating a clamp-like structure to encircle the DNA, allowing the extension of very distorting DNA damage (Fig.8A and Fig.9B).

**Figure 9.**
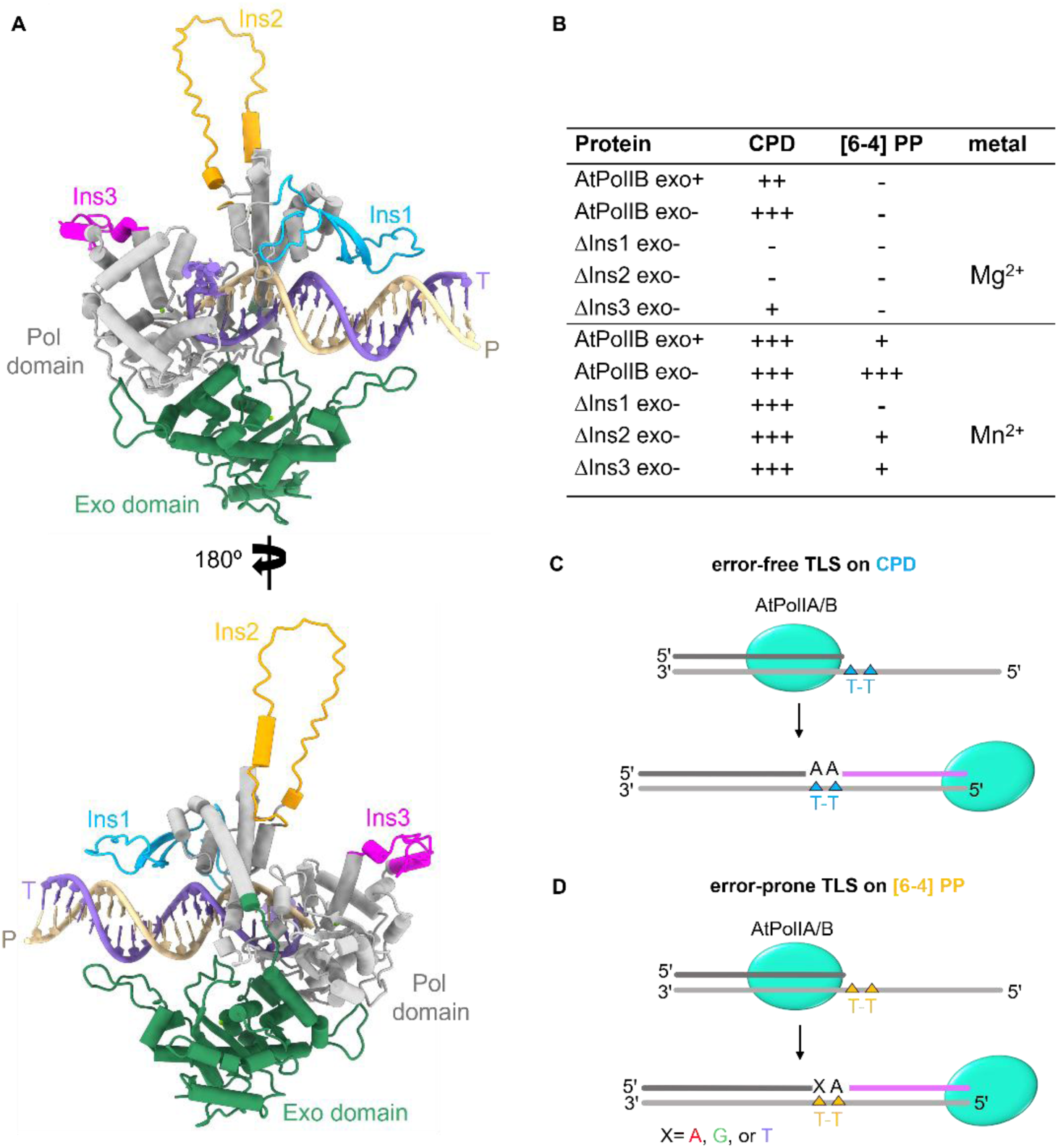
Proposed model of the Translesion DNA synthesis across UV lesions by plant organellar DNA polymerases. (**A**) AlphaFold prediction of AtPolIB (At3g20540) complexed with a primer/template DNA. Insertion1 is located from residues 576 to 621 (blue), insertion 2 mapped from residues 648 to 712 (orange), and insertion 3 from 844 to 869 (magenta). (**B**) Table indicating the TLS activity for each AtPolIB-derived variant in the presence of Mg^2+^ or Mn^2+^. TLS was ranked as limited (+), moderated (++) and strong (+++) TLS activity. (**C**) Cartoon indicating the error-free TLS activity by AtPolIs where preferentially incorporates adenines opposite CPD. (**D**) AtPolIA and AtPolIB insert A, G, T opposite the first thymine of [6-4] PP in an error-prone manner.

### 2. AtPolIs exhibit low exonucleolytic activity

Family-A DNAPs with an abrogated exonuclease active site perform TLS on CPDs, this would be due to the absence of primer shuttling into the exonuclease active site promoting multiple nucleotide insertion attempts (38, 55). Although AtPolls are replicative DNAPs, their 3’-5’ exonuclease or editing activity on dsDNA is lower than their counterpart in animals Pol γ (35, 56) (Fig. S3). This editing activity on AtPolIs also contrast to the potent 3’-5’ exonucleolytic activity of the 7DNAP-thioredoxin complex that is calculated to be near 10,000-folds more active (57). In contrast to T7DNAP that uses structural motifs to guide the 3’-end of a primer into the active site, AtPolIs lack those to direct primer shuttling from the polymerase into the exonuclease active site (58). Thus, lesion CPD lesion bypassing by AtPolls, specifically AtPoIIB correlates with the low exonucleolytic activity as a DNAP with strong editing activity as T7 DNAP can bypass CPD when the exonuclease activity is deleted (59). This hypothesis is confirmed when we tested the exonuclease-deficient AtPoIIs mutants which improved the ability to extend CPD and drastically enhanced DNA synthesis on the distorting [6-4] PP damaged (Fig. 6 compared to Fig. 3). Although AtPolIs exhibit a lower exonuclease activity respect the replicative human Pol γ, we discovered that AtPoIIs use error-free TLS on CPD while on [6-4] PP the bypassing mechanism became mutagenic (Fig. 7 and Fig. 9). Both AtPolIs preferentially added two Adenine opposite CPD and continue and efficient extension (Fig. 9A). Contrarily, during [6-4] PP synthesis both AtPolIs added A, G or T opposite the 3’-T of the dimer becoming a barrier for the next incorporation which results in the rate-limiting step due to formation of a T-G pair (Fig. 7F-I *right* and 9B). These observations explain why AtPolIs extend a [6-4] PP with very limited efficiency, and it is until the exonuclease domain is removed when TLS becomes stronger. TLS on [6-4] PP increased with the presence of Mn^2+^ suggesting that this metal would regulate the TLS activity by reshaping the catalytic site to accommodate very voluminous DNA damages.

### 3. AtPolIs have a strong binding affinity for the primer-template

Although the replicative T7 DNAP is unable to bypass CPD by itself, this enzyme bypasses CPD when forming a complex with the bacteriophage T7 helicase (60). TLS by this complex suggest that the intrinsic T7 DNAP TLS ability would be assisted by a protein that increases its affinity for the lesion-containing DNA. When we deleted Insertion 1, 2, 3 from AtPoIIB, these deletions abolished the TLS activity on CPD and [6-4] PP and while the presence of Mn^2+^ restored the ability on all deletion variants to bypass only CPD but did not restore the robust TLS on [6-4] PP as in AtPolIB exo-. Interestingly, ΔIns1 mutant also showed a lower extension band and intermediates products in reaction containing Mn^2+^ for the non-damaged substrate and not observed in the other variants, suggesting that insertion 1 would be involved in processivity and DNA binding (Fig. 8C-D). Numerous studies shown that the processivity factor PCNA increases the ability of replicative and TLS DNA polymerases to bypass DNA lesions (61–64). Our evidence suggests that AtPolIs TLS activity on UV lesions depends strongly on these three unique insertions in the polymerization domain (Fig. 8A) and resemble the feature found in Human Pol θ where similar insertions are implicated on primer-template stabilization during TLS activity opposite abasic sites and thymine glycols (65). Insertions variants only recovered the CDP TLS activity in the presence of Mn^2+^ which would suggest that the two intact insertions undergo structural re organizations important for a potential DNA encircling and increment of the DNA binding affinity.

### 4. Manganese ions modulate the flexibility of the polymerase active site

A recent study demonstrated that the CPD TLS activity of Human Pol γ, a family-A DNAP, would be triggered by Mn^2+^ (35). We found that both wild-type AtPolIs displayed TLS on CPD in the sole presence of Mg^2+^ while Mn^2+^ enhanced the extension and triggered a limited ability to bypass [6-4] PP damage. When we removed exonuclease activity on both AtPolIs, Mn^2+^ drastically increased TLS activity on [6-4] PP substrates.

Mg^2+^ and Mn^2+^ are essential micronutrients necessary for plant growth. The concentrations of free Mg^2+^ in chloroplast rank from 0.5 to 2.0 mM (68) and it can increase in a light-dependent manner up to 5 mM (66, 67), consistent with our results where the increment of Mg^2+^ facilitated the TLS activity on CPD substrates (Fig. 2). Whereas Mn^2+^ concentrations in plants rank between 0.1 to 1.5 mM while under stress conditions can reach up 2.0-2.5 mM (68, 69). This data indicates that in a given cellular condition AtPolls have both divalent metal ions available to be used, suggesting that the two metals can be present at similar concentrations but under certain scenarios the balance would change toward an increment of Mn^2+^ increasing the preference of AtPolIs at the extent that Mg^2+^ can be substituted for Mn^2+^ when DNA damages are encountered. Mn^2+^ may play a key role in reshaping critical regions in the protein such as the three unique insertions found in POPs but also allowing a more permissive catalytic pocket to ensure TLS on UV lesions.

All these together suggest that in the presence of Mg^2+^, the three insertions are important to bypass CPD and the absence of one of these abolished the activity, but the TLS would be restored if Mn^2+^ reshapes other critical protein regions. In contrast to CPD, [6-4] PP is a more voluminous and distorting UV lesion that may need the three intact insertions but also the presence of Mn^2+^ to reshape the catalytic pocket and additional regions (Fig. 9).

## SUPPLEMENTARY DATA STATEMENT

Supplementary Data are included in this file

## ACKNOWLEDGMENTS

We thank Dr. Rachael Ann DeTar for her insights regarding manganese and magnesium concentrations in chloroplasts.

## AUTHOR CONTRIBUTIONS STATEMENT

L.G.B., Y.W.Y. and N.B.T conceived the project. N.B.T. performed biochemical experiments, protein purification and data analysis. J.P. performed data analysis. E.C.T performed overexpression and protein purification. S.I. synthesized the templates containing UV lesion. L.G.B., Y.W.Y. and N.B.T. supervised the project. N.B.T. and L.G.B wrote the first draft, and all authors contributed to the final manuscript. L.G.B. and Y.W.Y. funding acquisition.

---

**Table S1.** Nucleotide incorporation by AtPolIA exo- using the N primer.

| % inserted nucleotide by AtPolIA<sub>exo</sub>- |  |  |  |  |  |  |  |  |  |  |  |  |
| --- | --- | --- | --- | --- | --- | --- | --- | --- | --- | --- | --- | --- |
|  | ND |  |  |  | CPD |  |  |  | [6-4] PP |  |  |  |
| template | A | T | G | C | A | T | G | C | A | T | G | C |
| A<sub>30</sub> | 7 | 0 | 0 | 0 | 0 | 0 | 0 | 0 | 0 | 0 | 0 | 0 |
| T<sub>29</sub> | 46 | 0 | 0 | 0 | 57 | 0 | 0 | 0 | 0 | 0 | 0 | 0 |
| T<sub>28</sub> | 0 | 0 | 0 | 0 | 0 | 6 | 16 | 0 | 86 | 15 | 30 | 0 |

**Table S2.** Nucleotide incorporation by AtPolIB exo- using the N primer.

| % inserted nucleotide by AtPolIB<sub>exo</sub>- |  |  |  |  |  |  |  |  |  |  |  |  |
| --- | --- | --- | --- | --- | --- | --- | --- | --- | --- | --- | --- | --- |
|  | ND |  |  |  | CPD |  |  |  | [6-4] PP |  |  |  |
| template | A | T | G | C | A | T | G | C | A | T | G | C |
| A<sub>30</sub> | 12 | 0 | 0 | 0 | 0 | 0 | 0 | 0 | 0 | 0 | 0 | 0 |
| T<sub>29</sub> | 44 | 0 | 0 | 0 | 66 | 0 | 0 | 0 | 0 | 0 | 0 | 0 |
| T<sub>28</sub> | 0 | 0 | 0 | 0 | 0 | 13 | 17 | 0 | 68 | 25 | 38 | 0 |

**Table S3.**
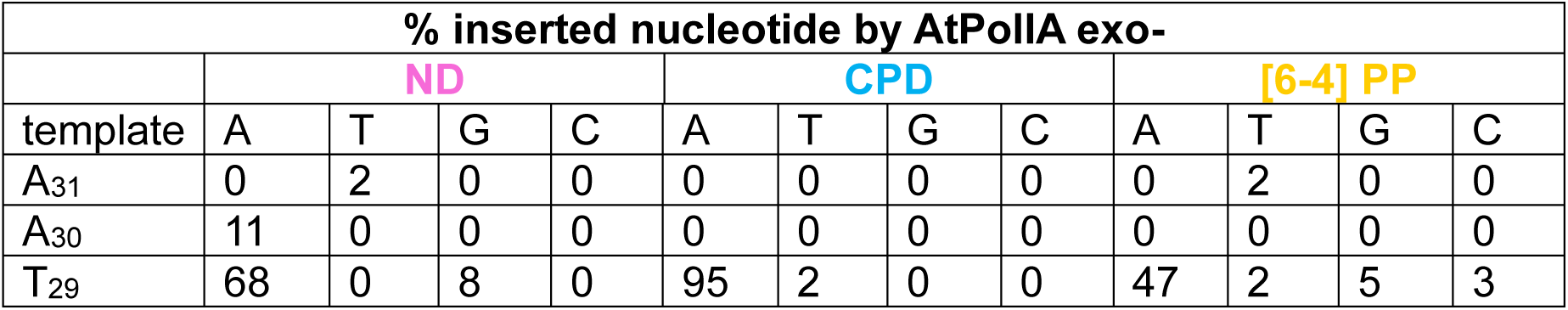
Nucleotide incorporation by AtPolIA exo- using the N+1 primer.

**Table S4.** Nucleotide incorporation by AtPolIB exo- using the N+1 primer.

| % inserted nucleotide by AtPolIB<sub>exo</sub>- |  |  |  |  |  |  |  |  |  |  |  |  |
| --- | --- | --- | --- | --- | --- | --- | --- | --- | --- | --- | --- | --- |
|  | ND |  |  |  | CPD |  |  |  | [6-4] PP |  |  |  |
| template | A | T | G | C | A | T | G | C | A | T | G | C |
| A<sub>31</sub> | 0 | 1 | 0 | 0 | 0 | 2 | 0 | 0 | 0 | 3 | 0 | 0 |
| A<sub>30</sub> | 3 | 0 | 0 | 0 | 0 | 0 | 0 | 0 | 0 | 0 | 0 | 0 |
| T<sub>29</sub> | 51 | 0 | 2 | 0 | 62 | 3 | 0 | 0 | 28 | 2 | 2 | 3 |

**Table S5.**
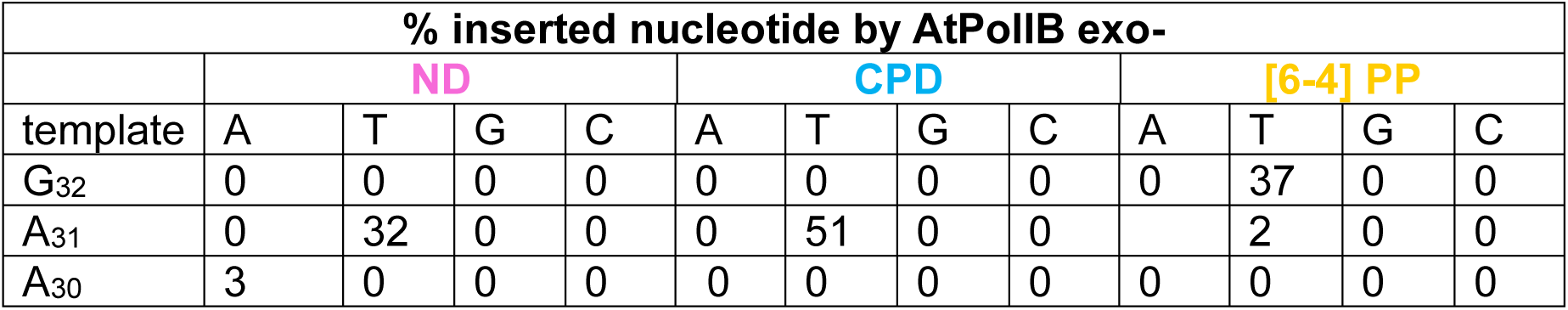
Nucleotide incorporation by AtPolIA exo- using the N+2 primer.

**Table S6.**
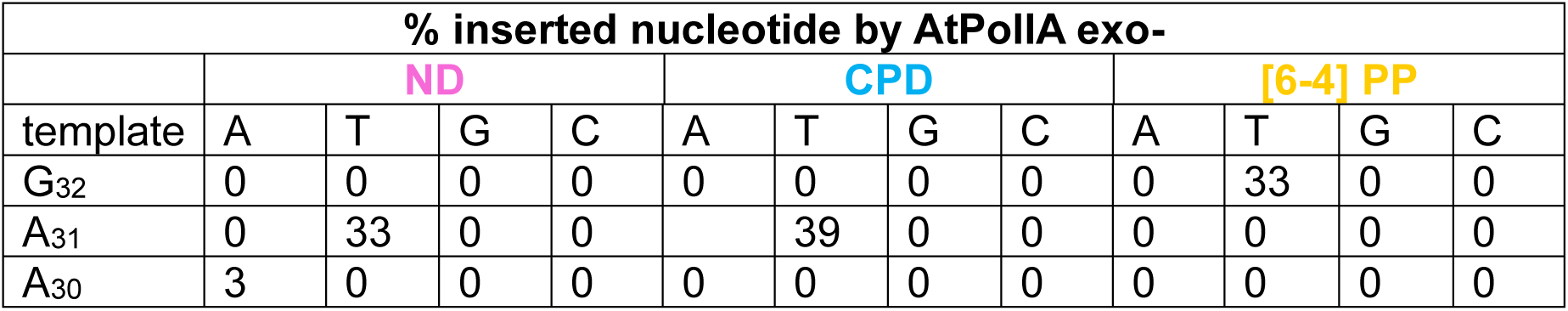
Nucleotide incorporation by AtPolIB exo- using the N+2 primer.

**Table S7.**
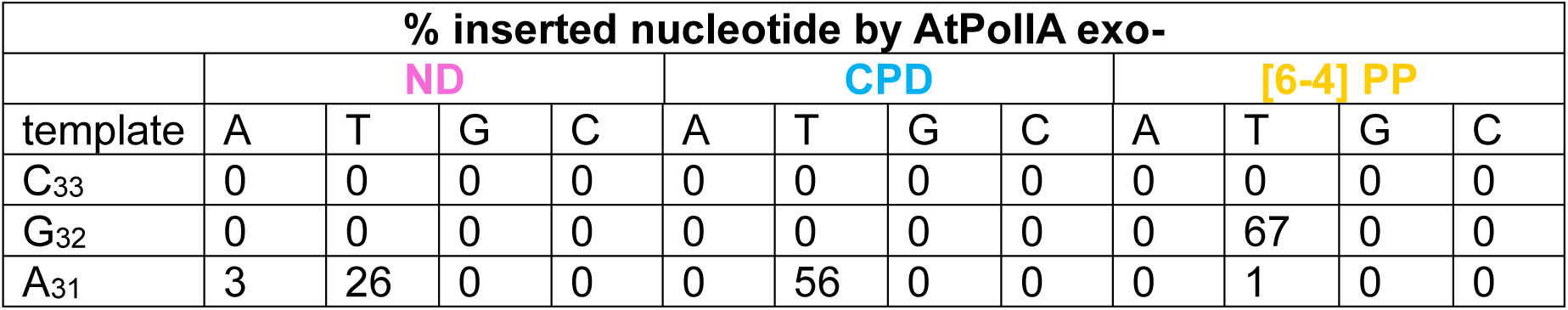
Nucleotide incorporation by AtPolIA exo- using the N+3 primer.

**Table S8.** Nucleotide incorporation by AtPolIB exo- using the N+3 primer.

| % inserted nucleotide by AtPolIIB exo- |  |  |  |  |  |  |  |  |  |  |  |  |
| --- | --- | --- | --- | --- | --- | --- | --- | --- | --- | --- | --- | --- |
|  | ND |  |  |  | CPD |  |  |  | [6-4] PP |  |  |  |
| template | A | T | G | C | A | T | G | C | A | T | G | C |
| C<sub>33</sub> | 0 | 0 | 0 | 0 | 0 | 0 | 0 | 0 | 0 | 0 | 0 | 0 |
| G<sub>32</sub> | 0 | 0 | 0 | 0 | 0 | 0 | 0 | 0 | 0 | 58 | 0 | 0 |
| A<sub>31</sub> | 3 | 49 | 0 | 0 | 2 | 41 | 0 | 2 | 0 | 4 | 0 | 0 |

**Figure S1.**
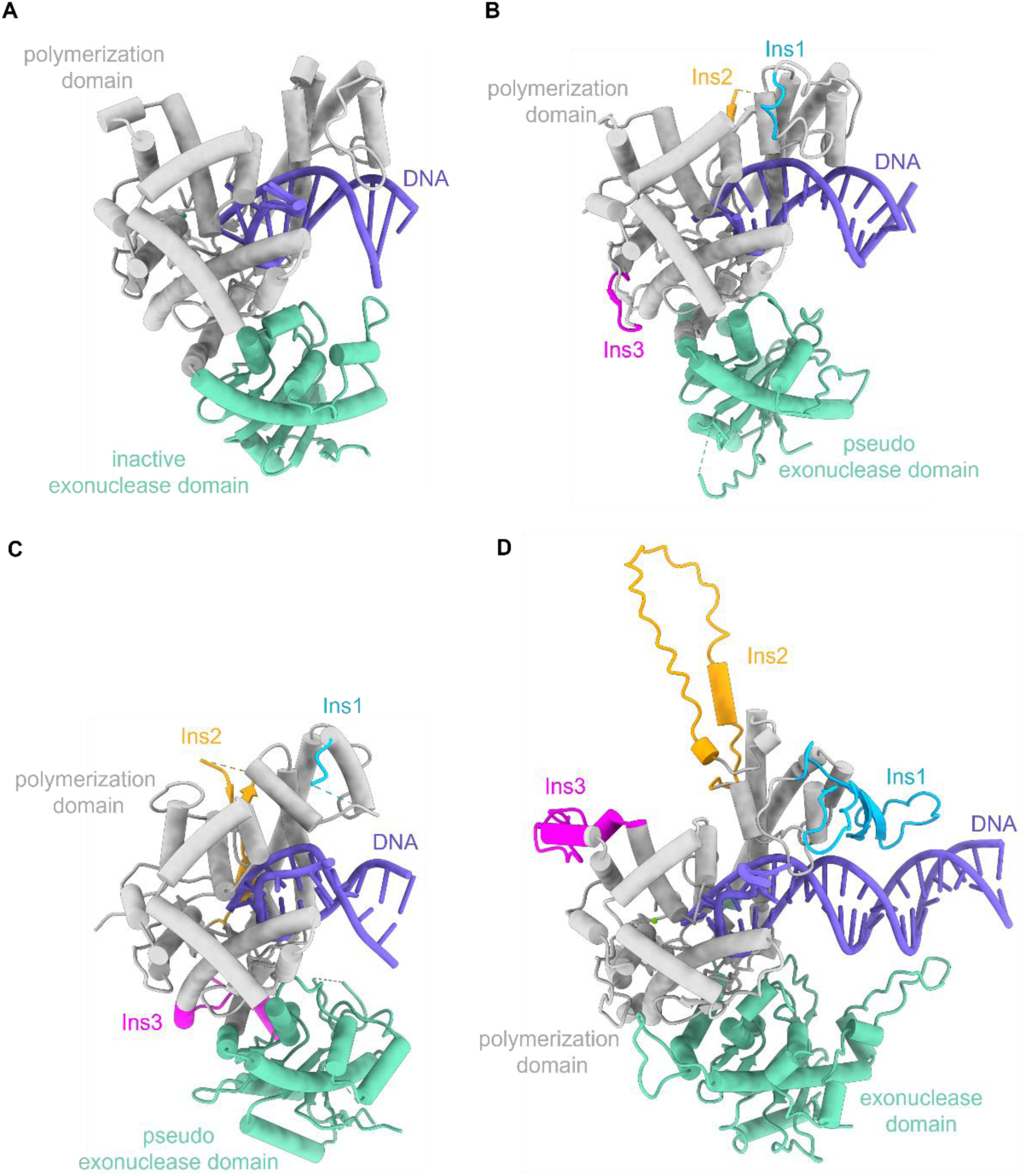
Structural comparison of AtPolIs with its evolutionary related family-A DNAPs. Crystal structures of *Bs* Pol, HsPol ν, and HsPol θ (45, 50, 68) in comparison with an Alpha-Fold prediction of AtPolIB in complex with dsDNA. (**A**) *Bs* Pol display its inactive exonuclease domain and polymerization domain but does not present similar insertion as AtPolIs. HsPol ν (**B**) and HsPol θ (**C**) display three insertions with Insertion1 close to the upstream duplex DNA and a pseudo exonuclease inactive domain. (**D**) AtPolIB contains three insertions: (Ins) 1, 2, and 3 corresponding to residues 575 to 623, 648 to 696, and 837 to 870, respectively. Insertion 1 is close to the upstream DNA creating an extra region for increasing DNA stability while insertion 2 and 3 are not directly in contact with the DNA. Insertions1, 2, and 3 are colored in blue, orange and magenta, respectively.

**Figure S2.**
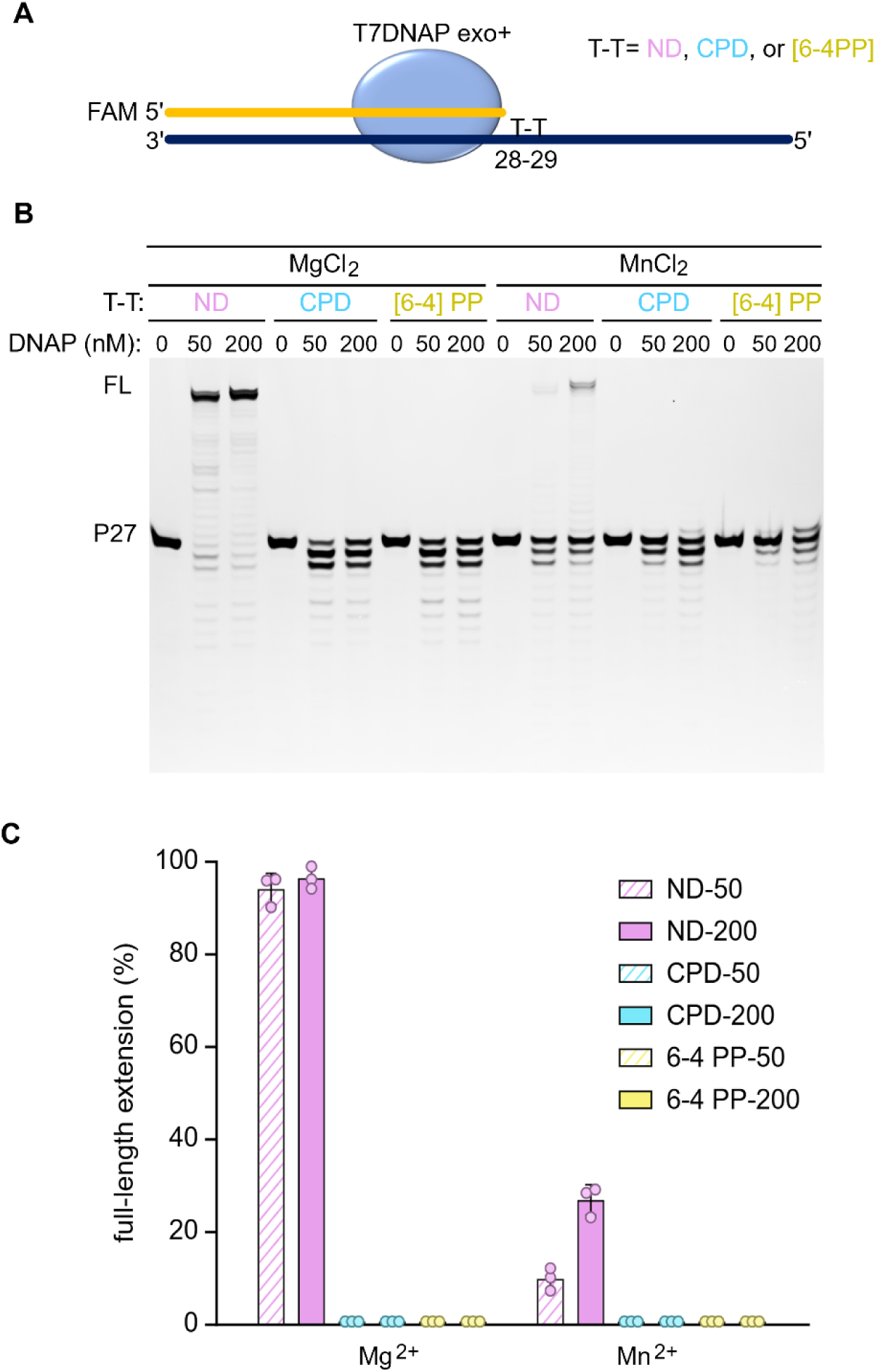
Primer extension assays by T7 DNAP exo+. (**A**) Scheme of the ND, CPD and [6-4] PP substrates used with a FAM labeled 27-nt primer right before the lesion. (**B**) Representative gel of the primer extension assay where 100 nM of DNA is incubated to 50 or 200 nM of T7DNAP exo+ in the presence of 10 mM of Mg^2+^ (*left panel*) or Mn^2+^ (*right panel*). Robust extension activity is observed in ND samples in the presence of Mg^2+^ and a reduced synthesis when Mn^2+^ is present. (**C**) Bar chart represents the percentage of extended primer for Non damaged (magenta), CPD (cyan) and [6-4] PP (yellow) from panel B. Error bars represent mean and standard deviation from at least two or three repeats.

**Figure S3.**
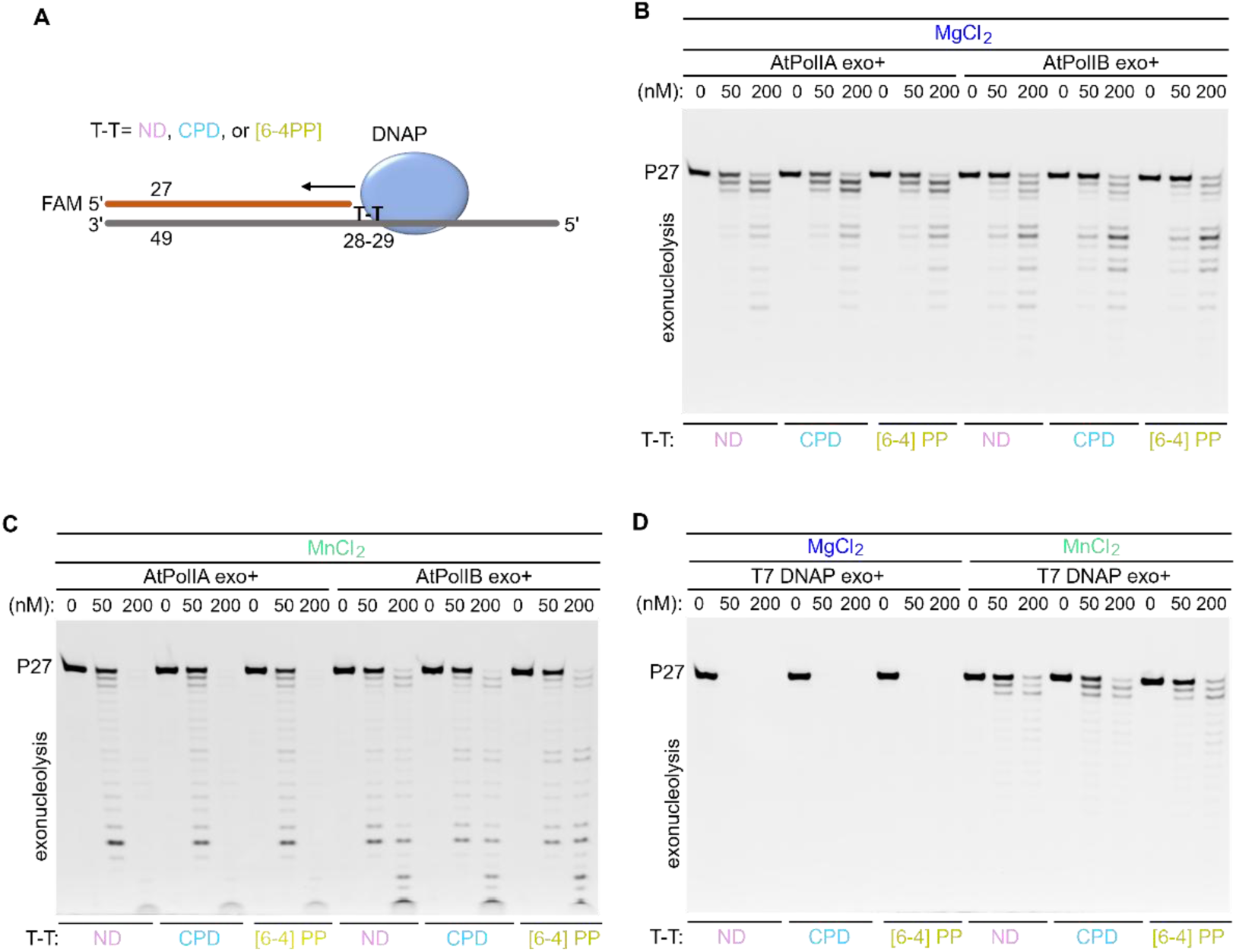
Exonuclease activity in the presence of Mg^2+^ or Mn^2+^ by AtPolIs. (**A**) Cartoon of the DNA templates containing a CPD or [6-4] PP lesion region after the 3’-OH end of FAM labeled primer. (**B**) 50 or 200 nM of AtPolIA exo+ (*left panel*) or AtPolIB exo+ (*right panel*) were incubated to 100 nM of DNA in the presence of 2 mM Mg^2+^. Both AtPolIs exhibited a similar exonucleolytic efficiency. (**C**) as in panel B but using 2 mM Mn^2+^. AtPolIA exo+ displays a higher exonuclease ability than AtPolIB exo+ at 200 nM (**D**) exonuclease assay by T7 DNAP exo+ in the presence of 10 mM of Mg^2+^ (*left panel*) or Mn^2+^ (*right panel*). Incubation with Mg^2+^ is very robust that exonucleolytic products run off the gel while the presence of Mn^2+^ reduces the exonuclease ability.

**Figure S4.**
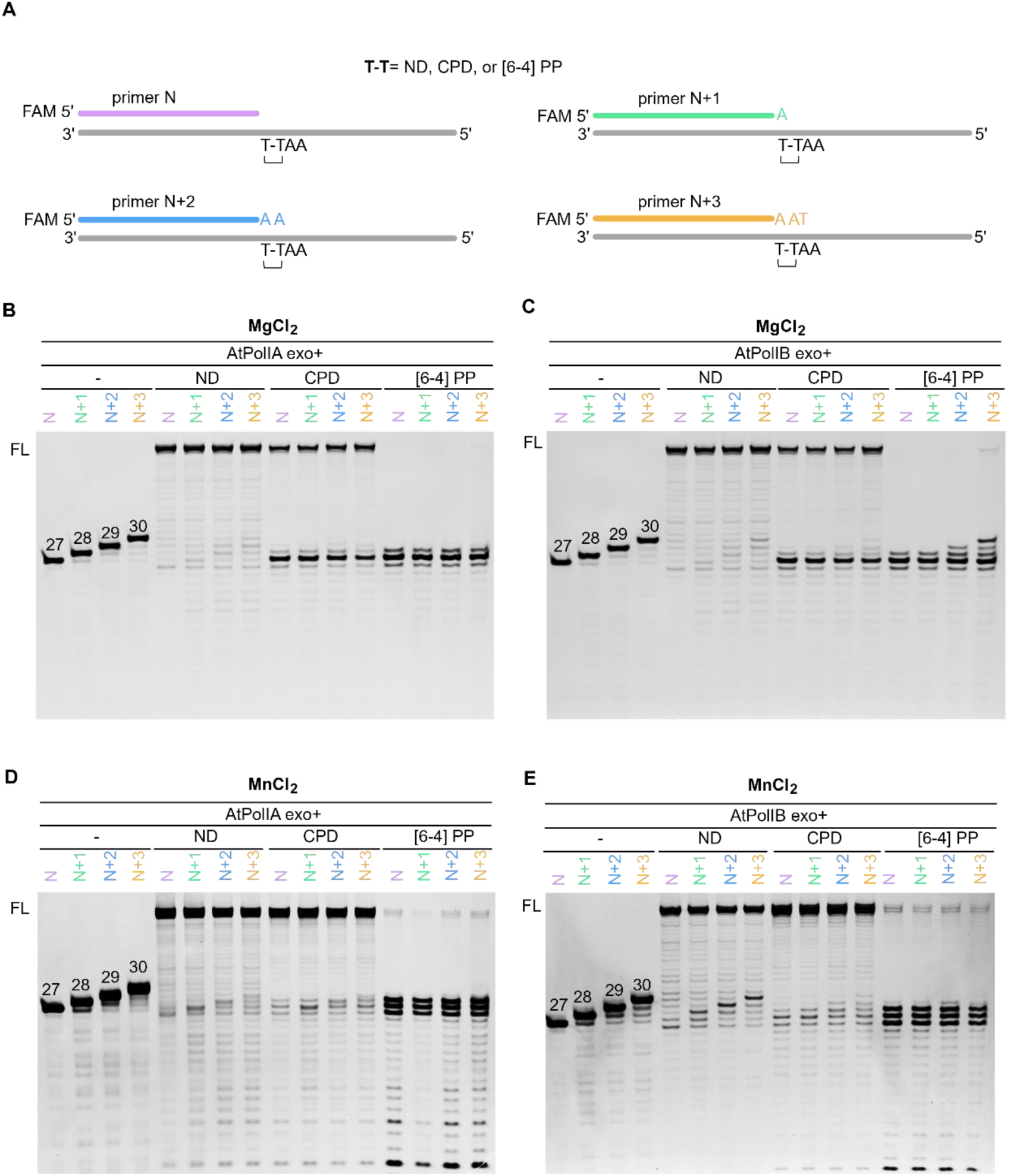
Lesion bypassing assay in the presence of Mg^2+^ or Mn^2+^ by exo-proficient AtPolIs. (**A**) Scheme of the four DNA substrates used for each template (ND, CPD or [6-4] PP). FAM labeled primer varies by 1 base: 27 (N, magenta), 28 (N+1, green), 29 (cyan) and 30 (N+3, yellow). T-T denotes the lesion or two thymine. (**B-E**) Representative gels from primer extension assays containing 100 nM of ND, CPD or [6-4] PP substrate and 200 nM of AtPolIA exo+ or AtPolIB exo+ in the presence of 2.0 mM Mg^2+^ (B-C) or Mn^2+^ (D-E). (-) primer control without enzyme.

**Figure S5.**
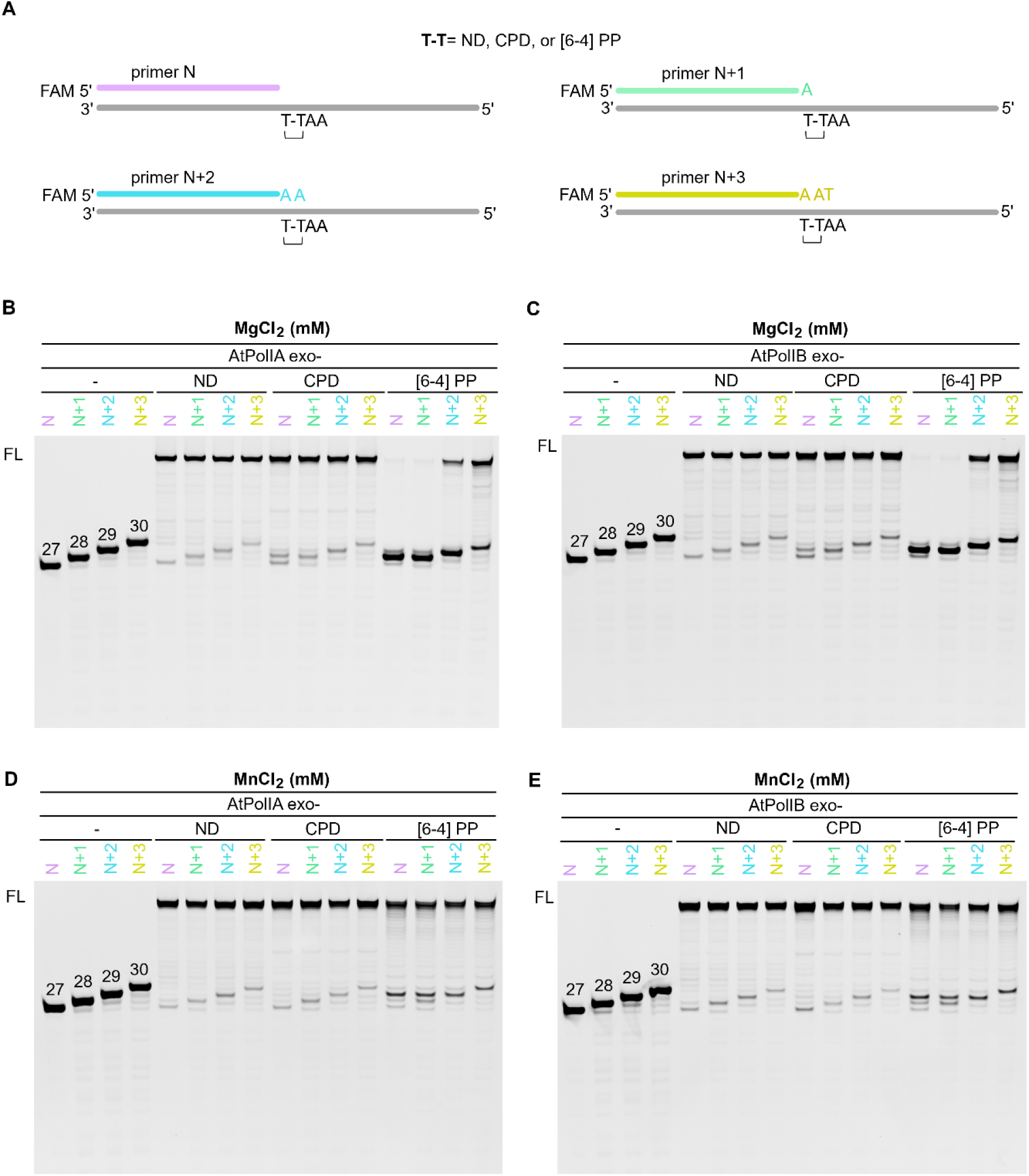
Lesion bypassing assay in the presence of Mg^2+^ or Mn^2+^ by exo-deficient AtPolIs. (**A**) Scheme of the four DNA substrates used for each template (ND, CPD or [6-4] PP). FAM labeled primer varies by 1 base: 27 (N, magenta), 28 (N+1, green), 29 (cyan) and 30 (N+3, yellow). T-T denotes the lesion or two thymine. (**B-E**) Representative gels from primer extension assays containing 100 nM of ND, CPD or [6-4] PP substrate and 200 nM of AtPolIA exo+ or AtPolIB exo+ in the presence of 2.0 mM Mg^2+^ (B-C) or Mn^2+^ (D-E). (-) primer control without enzyme.

